# Glucokinase links metabolism and lineage plasticity in neuroendocrine prostate cancer via interaction with AKT1

**DOI:** 10.64898/2026.01.06.697917

**Authors:** Kai Shen, Ruopeng Su, Yiyi Ji, Weiwei Zhang, Xinyu Liu, Xinyu Chai, Junyi Wang, Bo Liu, Ang Li, Haotian Wu, Tianxiang Wang, Xiang Zhou, Zhou Jiang, Helen He Zhu, Liang Dong, Yinjie Zhu, Baijun Dong, Jiahua Pan, Qi Wang, Wei Xue

**Author notes:** Corresponding author: Qi Wang, Department of Urology, Ren Ji Hospital, Shanghai Jiao Tong University School of Medicine, Shanghai 200120, China., Wei Xue, Department of Urology, Ren Ji Hospital, Shanghai Jiao Tong University School of Medicine, Shanghai200120. China. These authors contributed equally: Kai Shen, Ruopeng Su, and Yiyi Ji.

## Abstract

Cancer cells frequently rewire their metabolism to sustain growth and survival under stress. Despite the critical role of metabolic adaptation in tumorigenesis, how specific metabolic enzymes regulate lineage plasticity remains unclear. Here, through FDG-PET imaging and transcriptomic analyses, we reveal markedly elevated glucose uptake in neuroendocrine prostate cancer (NEPC) and identify glucokinase (GCK) as a MYCN-induced metabolic enzyme. Beyond its metabolic role, GCK is indispensable for maintaining the neuroendocrine lineage of prostate cancer cells by establishing a functional circuit with AKT1 through reciprocal regulation—AKT1 binds and phosphorylates GCK at S373, whereas GCK phosphorylates AKT1 at S473. Pharmacological disruption of this AKT1-GCK axis suppresses tumor growth in NEPC mouse models and patient-derived xenografts. Altogether, our findings uncover both the metabolic and noncanonical kinase functions of GCK and establish the AKT1-GCK axis as a key link between metabolic reprogramming and neuroendocrine lineage transition in prostate cancer.

## Introduction

Glucose provides the primary energy source for rapidly proliferating tumors, a circumstance that endows a survival advantage for cells who acquire extraordinary capability for glucose uptake. Over the past few decades, extensive research have focused on how cancer cells upregulate key enzymes and pathways to enhance glucose utilization^1^ and how this metabolic reprogramming can be exploited to visualize tumors in vivo using the radiolabeled glucose analogue [^18^F]-fluoro-2-deoxyglucose (FDG)^2,3^. This appreciation of glucose metabolism in cancer biology coincides with the recognition that major oncogenic drivers, including the PI3K-AKT-mTOR pathway, are frequently activated and influence metabolic activities^4,5^. Reciprocally, metabolic reprogramming extends beyond energy production, directly shaping cell fate through the diverse functions of metabolic enzymes that act as regulators of essential cellular processes^6^.

One such enzyme family is the hexokinases, which mediate the first committed step of glucose metabolism by catalyzing the glucose to glucose-6-phosphate, a central metabolite that connects glycolysis, pentose phosphate pathway, glycogenesis, and hexosamine biosynthesis. The hexokinase family comprises five major isoforms in mammals: HK1, HK2, HK3, HK4 (also known as glucokinase, GCK), and HKDC1. Previous studies, including ours, have demonstrated the critical roles of HK2 in tumor initiation and progression^7,8^. In contrast, research on GCK has largely focused on its role in glucose homeostasis and diabetes. While GCK was initially recognized as a glucose sensor in hepatocytes and pancreatic β cells^9^, recent evidence reveals broader physiological functions extending beyond its classical role in glucose regulation. In the central nervous system, neuronal GCK regulates glucose sensing and feeding behavior^10^; in regulatory T cells, GCK-driven glycolysis supports migration and immune homeostasis^11^; and in cardiomyocytes, activation of GCK by Hmbox1 inhibition enhances survival during ischemia/reperfusion injury^12^, suggesting a cytoprotective role under metabolic stress. Together, these findings indicate that GCK acts as a multifaceted regulator of systemic energy balance and stress adaptation, hinting at potential roles in cancer metabolism that remain unexplored.

Neuroendocrine prostate cancer (NEPC) is an aggressive subtype of prostate cancer characterized by small-cell morphology and elevated neuroendocrine markers, including synaptophysin (SYP), chromogranin A (CHGA), and CD56 (NCAM1)^13,14^. Genomic analyses reveal that NEPC frequently harbors loss of *PTEN*, *TP53*, and *RB1*, and combined deletion of these genes in mouse prostate epithelia induces tumors with neuroendocrine features, providing a robust preclinical model^15–17^. Moreover, MYCN overexpression and AKT1 activation can transform human prostate epithelial cells toward a neuroendocrine phenotype, underscoring their crucial role in lineage plasticity^18^. Although lineage-specific transcriptional programs in NEPC have been extensively studied, the accompanying metabolic adaptations that sustain this lineage switch remain poorly understood. Our previous work revealed high FDG uptake in patients with small-cell NEPC^19^, consistent with reports showing that FDG-PET outperforms PSMA-based imaging in detecting NEPC lesions^20,21^. Additionally, a neuroendocrine-related transcriptional signature positively correlates with the expression of glucose transporters and hexokinases^22^, indicating that reprogrammed glucose metabolism underlies the aggressive behavior of NEPC. These findings prompted us to investigate whether metabolic rewiring contributes to lineage plasticity and disease progression in NEPC.

In this study, we identify GCK as a MYCN-induced metabolic enzyme that is markedly elevated in NEPC and essential for maintaining the neuroendocrine lineage. Beyond its metabolic function, GCK interacts directly with AKT1, forming a reciprocal phosphorylation loop—in which AKT1 phosphorylates GCK at S373, while GCK phosphorylates AKT1 at S473. Pharmacological disruption of this GCK-AKT1 axis markedly impairs tumor growth in NEPC mouse models and patient-derived xenografts. Collectively, our findings reveal a noncanonical kinase function of GCK and define the GCK-AKT1 axis as a pivotal link that coordinates glucose metabolism and lineage transition in prostate cancer.

## Results

### NEPC shows high glucose uptake and elevated GCK expression

To characterize the changes of glucose uptake in prostate cancer, we retrospectively analyzed ^18^F-FDG PET/CT data from 416 patients with pathologically proven prostate cancer from Ren Ji Hospital (Table S1), including 41 diagnosed as NEPC and 375 as adenocarcinoma. As shown in Figure 1A, NEPC tumors showed significantly higher SUVmax values than adenocarcinoma, indicating enhanced intra-tumoral glucose uptake. We next imaged the ^18^F-FDG uptake in established genetically engineered mouse models, including *Pten* and *Trp53* double-knockout mice (Pb-Cre4: *Pten*^fl/fl^; *Trp53*^fl/fl^, DKO), which develop prostate adenocarcinoma, and *Pten*, *Trp53,* and *Rb1* triple-knockout mice (Pb-Cre4: *Pten*^fl/fl^; *Trp53*^fl/fl^; *Rb1*^fl/fl^, TKO), which recapitulate NEPC model. Consistent with the human data, tumors from TKO mice showed higher SUVmax value than those from DKO mice (Figure 1B). To further validate these findings in vitro, we compared glucose uptake among prostate cancer cells and found substantially higher uptake in NEPC cells LASCPC-01 than in adenocarcinoma cells (Figure 1C). We also assessed glucose uptake in induced neuroendocrine transition models, including MYCN-overexpressing PC3 cells and LNCaP cells overexpressing AR combined with TP53 and RB1 knockdown^17,23^. Both models displayed increased glucose uptake following neuroendocrine differentiation (Figures S1A and S1B). Together, these results indicate that NEPC lineage switching is accompanied by elevated glucose uptake.

**Figure 1.**
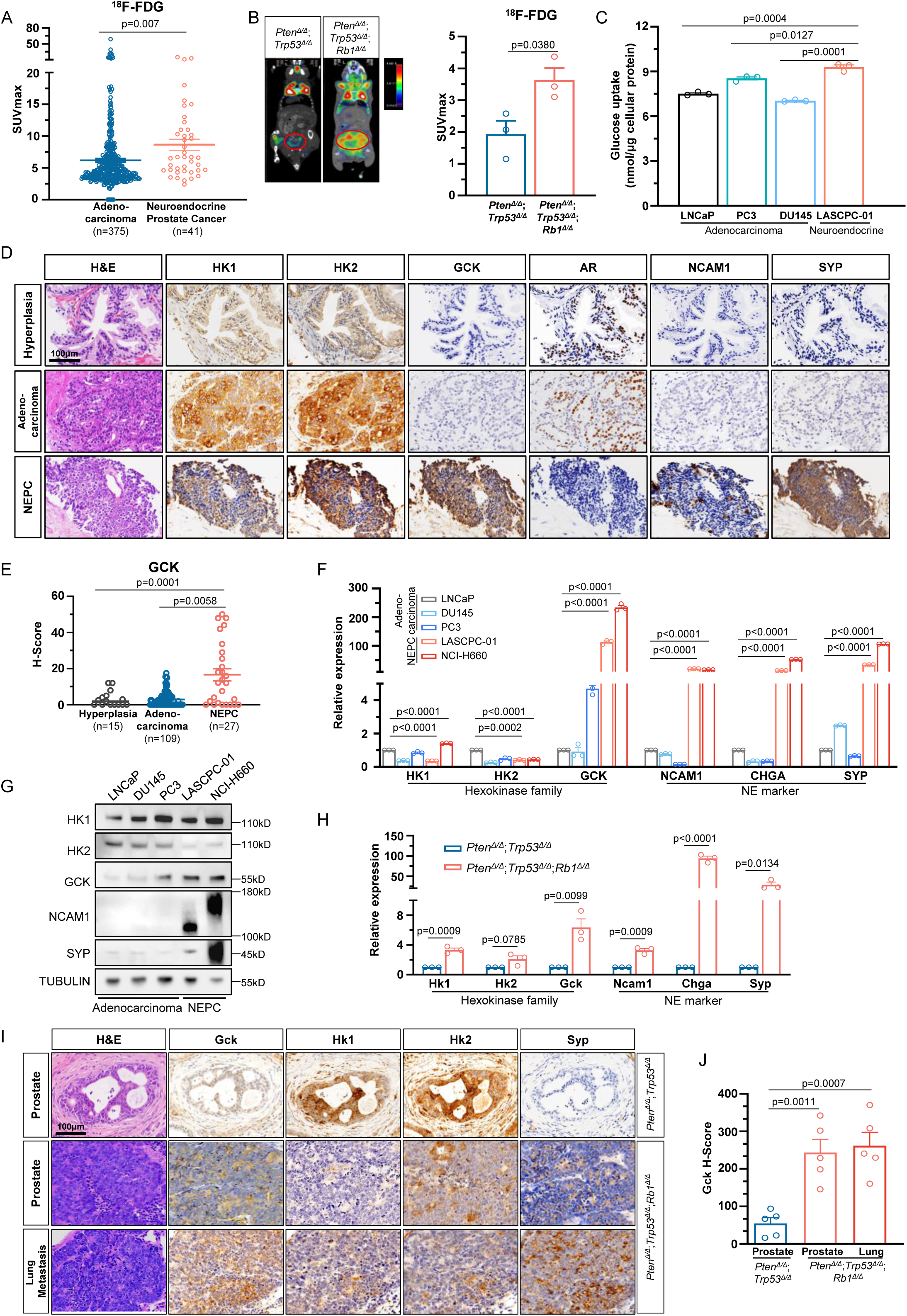
NEPC shows high glucose uptake and elevated GCK expression. (A) Scatter plot of SUVmax measured by ^18^F-FDG PET/CT for patients with prostate adenocarcinoma (n = 375) and neuroendocrine prostate cancer (n = 41). (B) Representative image (left) and SUVmax quantification (right) of ^18^F-FDG PET/CT in Pb-Cre4: *Pten*^fl/fl^; *Trp53*^fl/fl^ and Pb-Cre4: *Pten*^fl/fl^; *Trp53*^fl/fl^; *Rb1*^fl/fl^ mice (n = 3 for each group). (C) Glucose uptake assay in prostate adenocarcinoma and neuroendocrine prostate cancer cell lines (n = 3 biologically independent experiments). (D) Representative H&E and immunohistochemistry staining of hexokinase family, AR, NCAM1, and SYP in prostate samples from indicated groups (hyperplasia (n = 15), adenocarcinoma (n = 109), and NEPC (n = 27)). Scale bar, 100 μm. (E) Quantification of GCK staining in prostate samples from indicated groups, each dot represents sample from an individual patient. (F) QPCR showing relative mRNA expression of hexokinase family (*HK1*, *HK2*, and *GCK*) and neuroendocrine-related genes (*NCAM1*, *CHGA*, and *SYP*) in the indicated prostate cancer cells (n = 3 biologically independent experiments). (G) Western blot showing protein expression of hexokinase family (HK1, HK2, and GCK) and neuroendocrine-related protein (NCAM1 and SYP) in the indicated prostate cancer cells (n = 3 biologically independent experiments). (H) QPCR showing relative mRNA expression of hexokinase family (*Hk1*, *Hk2*, and *Gck*) and neuroendocrine-related genes (*Ncam1*, *Chga*, and *Syp*) in tumors from Pb-Cre4: *Pten*^fl/fl^; *Trp53*^fl/fl^ (n = 3) and Pb-Cre4: *Pten*^fl/fl^; *Trp53*^fl/fl^; *Rb1*^fl/fl^ mice (n = 3). (I) Representative H&E and immunohistochemistry staining of hexokinase family (Hk1, Hk2, and Gck) and Syp in tumors from Pb-Cre4: *Pten*^fl/fl^; *Trp53*^fl/fl^ (n = 6) and Pb-Cre4: *Pten*^fl/fl^; *Trp53*^fl/fl^; *Rb1*^fl/fl^ mice (n = 6). Scale bar, 100 μm. (J) Quantification of hexokinase family staining in samples from indicated mice, each dot represents sample from an individual mouse. Data presented as mean ± s.e.m (A, B, C, E, F, H, J). Statistical significance was determined by two-tailed unpaired Student’s test (A, B, C, E, H, J) or one-way analysis of variance (ANOVA) with Dunnett’s multiple comparisons (F).

Given that intracellular glucose retention depends on phosphorylation by hexokinase family, which catalyzed the first committed step of glucose metabolism, we next examined the expression profiles of hexokinase family members in prostate cancer datasets encompassing both adenocarcinoma and NEPC. Unexpectedly, GCK, rather than HK2, showed the most pronounced upregulation in NEPC (Figures S1C and S1D). Consistent patterns were observed in mouse models, where Gck expression was markedly increased in NEPC tumors relative to wildtype or adenocarcinoma counterparts (Figures S1E and S1F). Analysis of patient bulk transcriptomic data showed that GCK, unlike other hexokinases, was correlated positively with neuroendocrine markers and negatively with AR-related genes (Figure S1G). Single-cell RNA sequencing data from six castration-resistant prostate cancer patients, including two with histologically confirmed NEPC, further supported these findings. Unsupervised clustering of 14,210 epithelial cells identified two clusters enriched in neuroendocrine features. GCK expression was predominantly enriched in one of these clusters, consistent with its association with the neuroendocrine phenotype (Figure S1H). In line with the bulk RNA-seq data, GCK expression correlated positively with neuroendocrine-related and inversely with AR-related gene signatures (Figure S1I). To confirm these observations at the protein level, we performed immunohistochemical analysis on prostate tissue samples from 151 patients, including benign prostatic hyperplasia (n = 15), adenocarcinoma (n = 109), and NEPC (n = 27). Quantitative analysis revealed a striking increase in GCK staining intensity and percentage in NEPC tissues (Figures 1D and 1E). Similarly, Western blot and qPCR analyses demonstrated higher GCK expression in NEPC-associated cell lines (NCI-H660 and LASCPC-01) than in adenocarcinoma-derived lines (Figures 1F and 1G).

Finally, we confirmed the differential expression of GCK in mouse models. In adenocarcinoma (Pb-Cre4: *Pten*^fl/fl^; *Trp53*^fl/fl^) and NEPC (Pb-Cre4: *Pten*^fl/fl^; *Trp53*^fl/fl^; *Rb1*^fl/fl^) models, Hk2 expression remained largely unchanged, and Hk1 showed only a modest change, whereas Gck was dramatically upregulated in NEPC tumors at both mRNA and protein levels, a pattern that persisted in lung metastases (Figures 1H–1J). Together, these findings identify GCK as the predominantly elevated hexokinase isoform in NEPC and link its upregulation to the enhanced glucose uptake characteristic of this aggressive prostate cancer subtype.

### GCK is essential for maintaining the neuroendocrine lineage in prostate cancer

Given its marked elevation in NEPC, we next examined whether GCK might also influence the neuroendocrine lineage transition in prostate cancer cells. To test this possibility, we ectopically overexpressed GCK in adenocarcinoma cell lines LNCaP, PC3, and DU145. However, the expression of neuroendocrine markers remained unchanged or only modestly increased upon GCK overexpression, indicating that GCK alone is insufficient to induce neuroendocrine switch in prostate cancer cells (Figures S2A–S2E). In contrast, silencing GCK with shRNA dramatically reduced neuroendocrine features in NEPC cell lines LASCPC-01, LNCaP/AR/shTP53/shRB1, and NCI-H660 (Figures 2A, 2B, 2C, 2D, and S2F). Consistent results were obtained in an NEPC patient-derived organoid model, in which GCK depletion led to the loss of neuroendocrine characteristics (Figure 2E). We further validated these findings using CRISPR/Cas9-mediated GCK knockout in LASCPC-01 cells (Figures 2F and 2G), confirming the essential role of GCK in maintaining the neuroendocrine lineage. Next, we assessed the impact of GCK on cell proliferation. Knockdown or knockout of GCK significantly reduced cell viability in LASCPC-01, LNCaP/AR/shTP53/shRB1, and the patient-derived organoid (Figures 2H, 2I, 2J, and S2G). This inhibitory effect was further supported by colony formation assays, which showed decreased clonogenic potential in LNCaP/AR/shTP53/shRB1 cells with GCK knockdown (Figures S2H and S2I). This effect was even more pronounced in xenograft models using LASCPC-01 cells (Figure 2K). Immunohistochemical staining of xenograft tumors revealed that GCK knockdown markedly decreased neuroendocrine marker expression (SYP), suppressed proliferation (Ki-67), and induced apoptosis (TUNEL) (Figure 2L). Given that NEPC typically exhibits resistance to androgen-targeted therapy, we further tested whether GCK influences this phenotype. As shown in Figures 2M and 2N, GCK overexpression significantly increased the IC_50_ for bicalutamide and enzalutamide in LNCaP cells, suggesting enhanced drug resistance. Together, these results demonstrate that GCK is critical for sustaining neuroendocrine lineage identity, supporting tumor proliferation, and promoting resistance to androgen-targeted therapies in NEPC cells.

**Figure 2.**
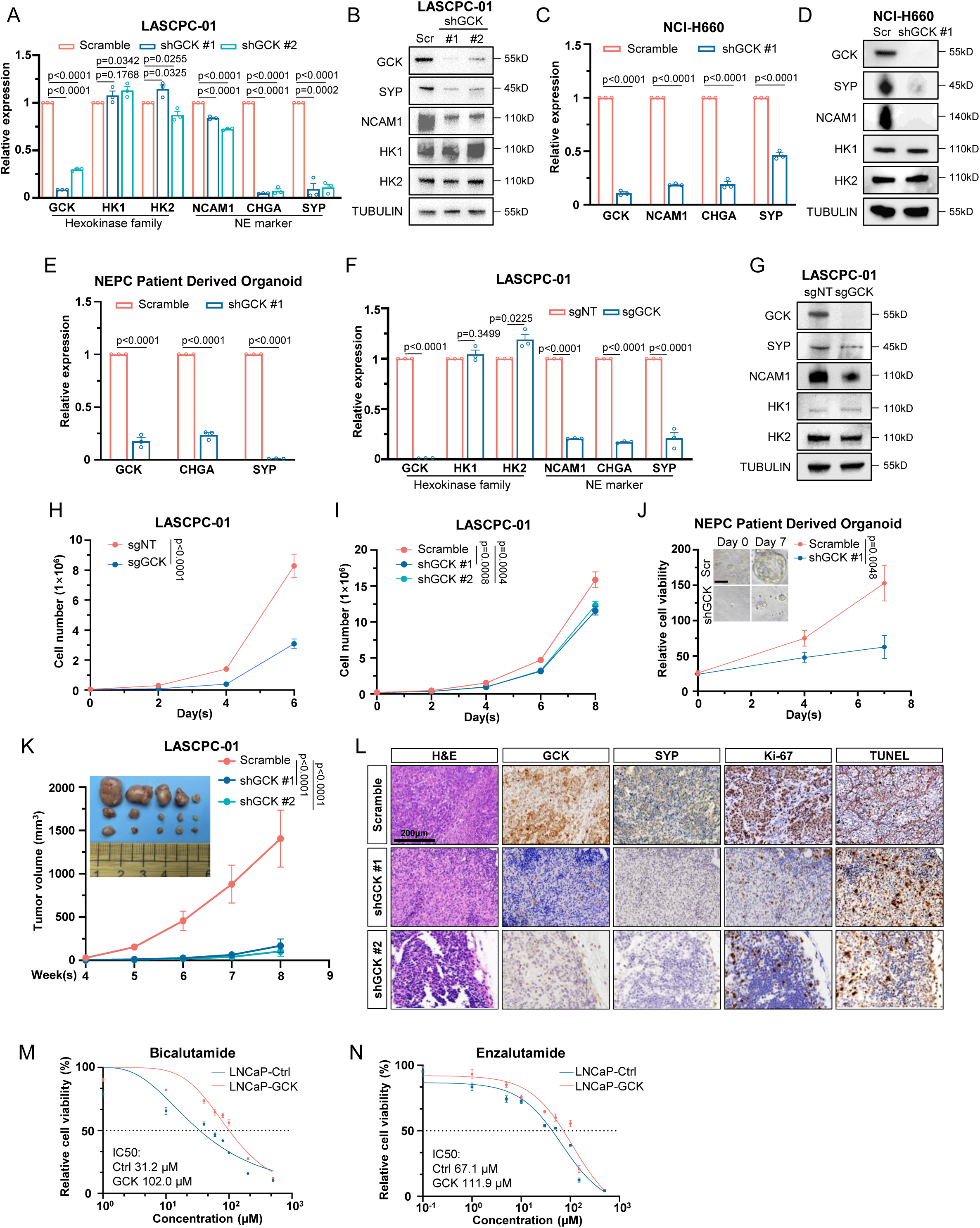
GCK is essential for maintaining the neuroendocrine lineage in prostate cancer. (A) QPCR showing relative mRNA expression genes of hexokinase family (*HK1*, *HK2*, and *GCK*) and neuroendocrine-related genes (*CHGA*, *NCAM1*, and *SYP*) in LASCPC-01/scramble, LASCPC-01/shGCK #1, and LASCPC-01/shGCK #2 (n = 3 biologically independent experiments). (B) Western blot showing indicated protein expression of hexokinase family (HK1, HK2, and GCK) and neuroendocrine-related proteins (NCAM1 and SYP) in LASCPC-01/scramble, LASCPC-01/shGCK #1, and LASCPC-01/shGCK #2 (n = 3 biologically independent experiments). (C) QPCR showing relative mRNA expression genes in NCI-H660/scramble and NCI-H660/shGCK #1 (n = 3 biologically independent experiments). (D) Western blot showing indicated protein expression in NCI-H660/scramble and NCI-H660/shGCK #1(n = 3 biologically independent experiments). (E) QPCR showing relative mRNA expression genes in PDO/scramble and PDO/shGCK #1 (n = 3 biologically independent experiments). (F) QPCR showing relative mRNA expression of indicated genes in LASCPC-01/sgNT and LASCPC-01/sgGCK (n = 3 biologically independent experiments). (G) Western blot showing indicated protein expression in LASCPC-01/sgNT and LASCPC-01/sgGCK. (H) Proliferation rate of LASCPC-01/sgNT and LASCPC-01/sgGCK cells at indicated time points (n = 5 biologically independent experiments). (I) Proliferation rate of LASCPC-01/scramble, LASCPC-01/shGCK #1, and LASCPC-01/shGCK #2 cells at indicated time points (n = 3 biologically independent experiments). (J) Cell viability of PDO/scramble and PDO/shGCK #1 cells at different time points. Photograph showing the represent image of organoids at day 0 and day 7 (n = 6 biologically independent experiments). Scale bar, 20 μm. (K) Photograph showing LASCPC-01 scramble, shGCK #1, and shGCK #2 xenografts in nude mice. Tumor volume was measured once a week at indicated time points (n = 5 per group). (L) Representative H&E and immunohistochemistry staining of GCK, SYP, Ki-67, and TUNEL in LASCPC-01 scramble, shGCK #1, and shGCK #2 xenografts. Scale bar, 200 μm. (M) Cell viability of LNCaP/Ctrl and LNCaP/GCK treated with indicated concentration of Bicalutamide for 48h (n = 3 biologically independent experiments). (N) Cell viability of LNCaP/Ctrl and LNCaP/GCK treated with indicated concentration of Enzalutamide for 48 h (n = 3 biologically independent experiments). Data presented as mean ± s.e.m (A, C, E, F, H, I, J, K, M, N). Statistical significance was determined by two-tailed unpaired Student’s test (A, C, E, F) or one-way analysis of variance (ANOVA) with Dunnett’s multiple comparisons (H, I, J, K, M, N).

### MYCN transcriptionally upregulates GCK

Next, we sought to identify the upstream regulatory pathways responsible for the upregulation of GCK expression in NEPC cells. Previous studies have shown that c-MYC, a member of the MYC family that shares a conserved DNA-binding domain with MYCN, can bind to the promoter of the HK2 gene and activate its transcription^24^. Given that MYCN serves as a key lineage-defining transcription factor in neuroendocrine tumors, and that MYCN expression shows a strong positive correlation with GCK levels in NEPC patients samples (Figures S1G and S1I), we hypothesized that MYCN might directly regulate GCK expression in NEPC. Indeed, c-MYC upregulated HK2 but had minimal effect on GCK (Figures S3A and S3B), whereas MYCN strongly induced GCK expression but only modest effects on HK2 (Figures 3A and 3B). By scanning the GCK promoter region, we identified three putative MYCN-binding sites (BS1, BS2, and BS3). Two promoter fragments encompassing these sites (Segment A and Segment B) were cloned into luciferase reporter plasmids (Figure 3C). Luciferase reporter assays revealed that MYCN significantly activated the transcriptional activity of both segments in HEK293T cells (Figure 3D). To determine the functional relevance of these binding sites, we mutated each CACGTG/CACCTG consensus sequence to CACATG, and found that disruption of any single site was sufficient to disrupt the binding of MYCN (Figure 3E). ChIP-qPCR analysis further validated the direct binding of MYCN to the *GCK* promoter (Figure 3F). Moreover, CRISPR/Cas9-mediated deletion of endogenous MYCN-binding sites in PC3 cells abolished MYCN-induced *GCK* transcription, confirming the specificity of this regulation (Figures 3G and S3C). In line with previous studies demonstrating that MYCN is sufficient to induce neuroendocrine switch in prostate cancer^18^, we found that GCK knockdown effectively abrogated MYCN-induced neuroendocrine marker expression (Figures 3H, 3I, and S3D) and suppressed MYCN-driven cell proliferation (Figures S3E–S3G). Collectively, these findings establish GCK as a direct transcriptional target of MYCN and reveal that GCK is indispensable for MYCN-mediated neuroendocrine lineage plasticity and cell proliferation in prostate cancer.

**Figure 3.**
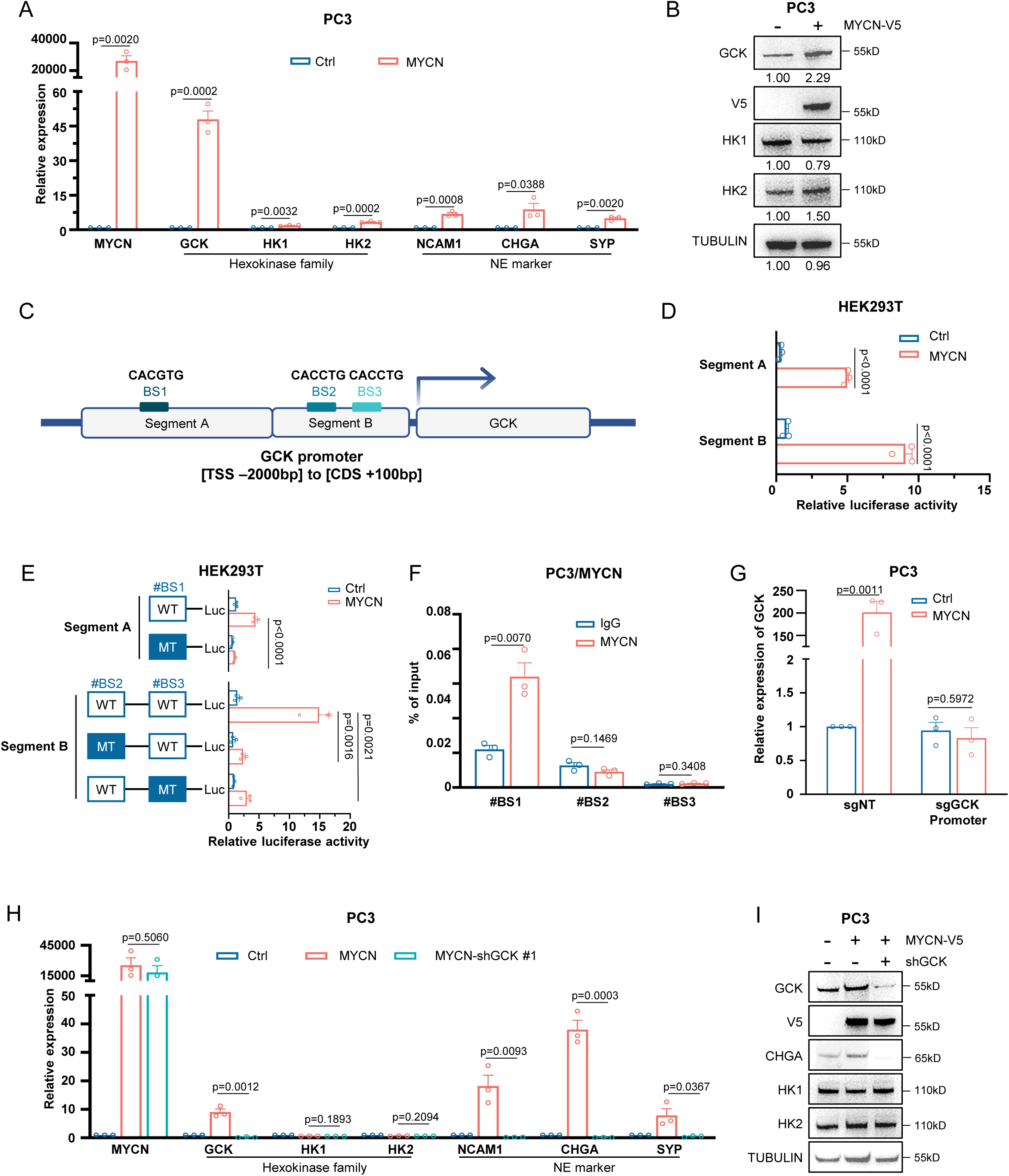
MYCN transcriptionally upregulates GCK. (A) QPCR showing relative mRNA expression of indicated genes in PC3/Ctrl and PC3/MYCN (n = 3 biologically independent experiments). (B) Western blot showing indicated protein expression in PC3/Ctrl and PC3/MYCN-V5 (n = 3 biologically independent experiments). (C) Schematic diagram showing putative MYCN binding motifs on *GCK* promoter and segments designed for luciferase reporter assay. (D) Relative luciferase activities driven by indicated segments of *GCK* promoter in HEK293T/Ctrl and HEK293T/MYCN (n = 3 biologically independent experiments). (E) Relative luciferase activities driven by wildtype or indicated mutant Segment of *GCK* promoter reporters in HEK293T/Ctrl and HEK293T/MYCN (n = 3 biologically independent experiments). (F) ChIP-qPCR of different locations on *GCK* promoter from PC3/MYCN immunoprecipitation with either anti-MYCN antibody or anti-IgG antibody (n = 3 biologically independent experiments). (G) QPCR showing relative *GCK* mRNA expression in PC3/sgNT and PC3/sgGCK promoter with/without the overexpression of MYCN. The sgRNAs were designed to target the MYCN binding sites in the *GCK* promoter region and the non-targeting sgRNA was used as control (n = 3 biologically independent experiments). (H) QPCR showing relative mRNA expression of indicated genes in PC3/Ctrl, PC3/MYCN, and PC3/MYCN/shGCK #1 (n = 3 biologically independent experiments). (I) Western blot showing indicated protein expression in PC3/Ctrl, PC3/MYCN, and PC3/MYCN/shGCK #1. Data presented as mean ± s.e.m (A, D, E, F, G, H). Statistical significance was determined by two-tailed unpaired Student’s test (A, D, E, F, G, H).

### The function of GCK in NEPC is independent of its metabolic activity

Given that GCK is transcriptionally activated by MYCN and essential for maintaining the neuroendocrine phenotype, we next sought to determine whether its function depends on its canonical enzymatic activity in glucose metabolism. GCK classically functions as a member of the hexokinase family, responsible for phosphorylating glucose and trapping it inside the cells, thereby promoting cellular glucose uptake. We thus asked whether the role of GCK in maintaining neuroendocrine plasticity depends on its glucose kinase activity. While ectopic expression of GCK in adenocarcinoma cells mildly increased glucose uptake (Figures 4A and 4B), knockdown of GCK in NEPC cell line LASCPC-01 strongly decreased glucose uptake (Figure 4C). We next introduced point mutations at two key amino acid residues, S151 and K169, within the hexokinase domain of GCK, which are critical for glucose binding^8,25,26^. As expected, these mutants displayed substantially reduced glucose kinase activity compared with wildtype GCK (Figures 4D and 4E). However, when we re-introduced shRNA-resistant wildtype or mutant GCK constructs into GCK-depleted LASCPC-01 cells, we found that the mutant forms, either singly or in combination, restored the reduced cell viability caused by GCK knockdown to a level comparable to wildtype GCK (Figure 4F). Similarly, GCK mutants were equally effective as the wildtype protein in rescuing the neuroendocrine phenotype in shGCK LASCPC-01 cells (Figures 4G and 4H). Altogether, these findings suggest that although GCK enhances glucose uptake in NEPC cells, its essential role in sustaining neuroendocrine lineage identity is independent of its canonical metabolic activity.

**Figure 4.**
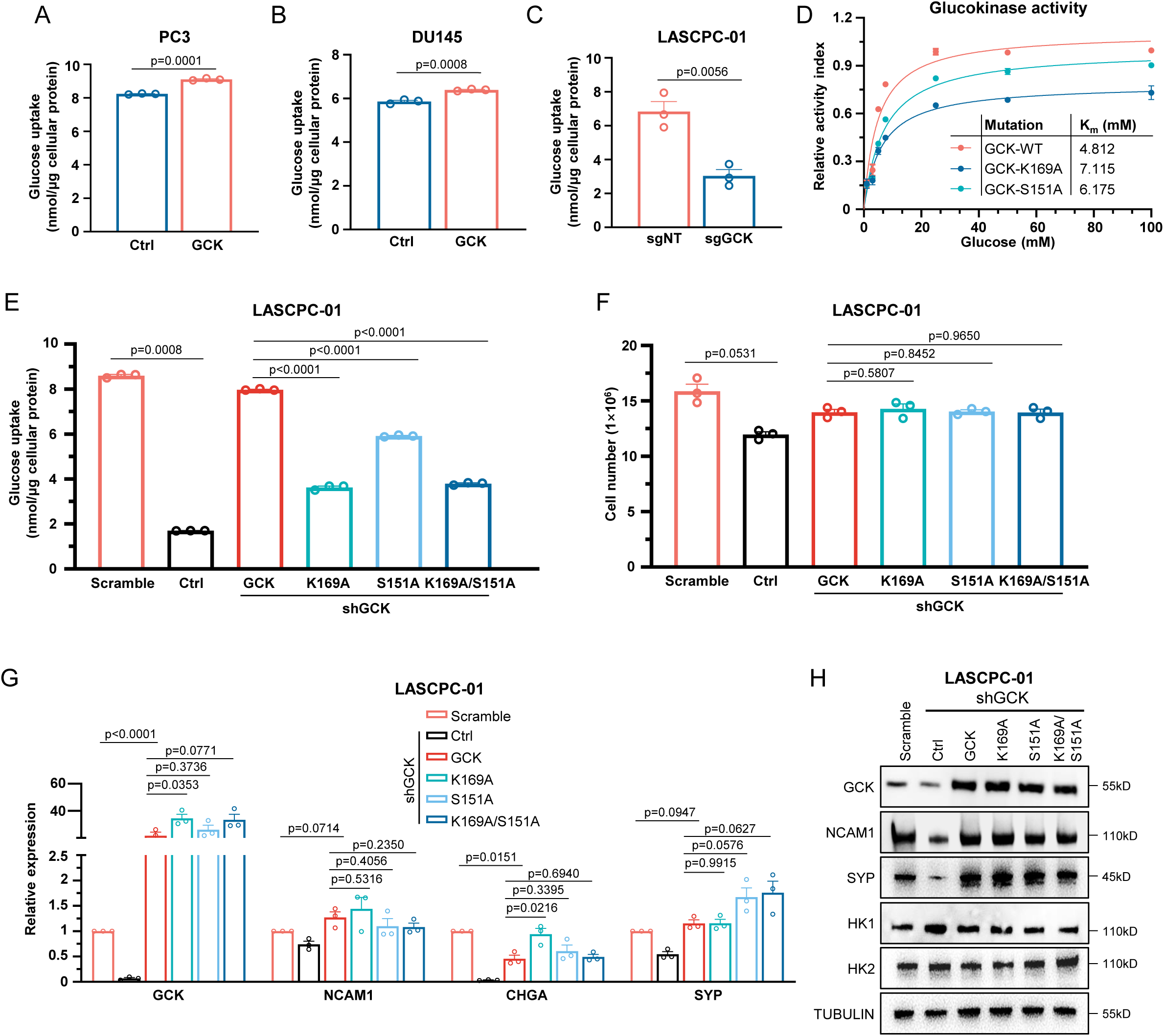
The function of GCK in NEPC is independent of its metabolic activity. (A) Glucose uptake assay in PC3/Ctrl and PC3/GCK (n = 3 biologically independent experiments). (B) Glucose uptake assay in DU145/Ctrl and DU145/GCK (n = 3 biologically independent experiments). (C) Glucose uptake assay in LASCPC-01/sgNT and LASCPC-01/sgGCK (n = 3 biologically independent experiments). (D) Glucokinase activity assay in wildtype GCK and indicated GCK mutations. GCK-V5 proteins were purified from HEK293T cell lysate with V5 antibody (n = 3 biologically independent experiments). (E) Glucose uptake assay in LASCPC-01 cells transfected with indicated GCK mutations (n = 3 biologically independent experiments). (F) Proliferation rate of indicated LASCPC-01 cells after 8 days of culture (n = 3 biologically independent experiments). (G) QPCR showing relative mRNA expression of indicated genes in LASCPC-01 cells transfected with indicated different GCK mutations (n = 3 biologically independent experiments). (H) Western blot showing indicated protein expression in LASCPC-01 cells transfected with indicated different GCK mutations (n = 3 biologically independent experiments). Data presented as mean ± s.e.m (A, B, C, D, E, F, G). Statistical significance was determined by two-tailed unpaired Student’s test (A, B, C, E, F) or one-way analysis of variance (ANOVA) with Dunnett’s multiple comparisons (D, G).

### GCK promotes lipid metabolism in NEPC

Having demonstrated that the lineage-sustaining function of GCK is independent of its canonical glucose-phosphorylating activity, we next sought to elucidate its molecular functions in NEPC cells. To this end, we performed RNA sequencing on LASCPC-01 cells with shRNA-mediated GCK knockdown and scramble control. As shown in Figure 5A, a total of 903 genes were upregulated and 997 genes were downregulated upon GCK depletion. Consistent with earlier findings, GCK deficiency was negatively correlated with neuroendocrine-associated genes and positively correlated with androgen receptor-related genes (Figure S4A). Gene Set Enrichment Analysis (GSEA) revealed a significant negative enrichment of the NEPC signature^27^, while KEGG pathway analysis indicated upregulation of apoptosis-related pathways and downregulation of neuronal differentiation-related pathways upon GCK knockdown (Figures 5B and 5C). Together, these data reinforce the essential role of GCK in sustaining neuroendocrine lineage identity and cell proliferation. Unexpectedly, we also observed a pronounced negative enrichment of multiple lipid metabolism-related pathways in GCK-deficient cells, including lipid homeostasis, sterol homeostasis, fatty acid homeostasis, phospholipid homeostasis, positive regulation of lipid catabolic process, and cholesterol homeostasis (Figure 5C). Transcriptome analysis further showed that GCK depletion downregulated key genes involved in lipid biosynthesis, particularly those in fatty acid synthesis (*ACLY*, *FASN*, *ACSL1*, and *SREBP1*) and cholesterol synthesis (*ACAT2*, *HMGCR*, and *SREBP2*) (Figure S4A), suggesting a close association between GCK expression and lipid metabolism. To validate this observation, we examined the expression of lipid metabolism-related genes in LASCPC-01 cells and confirmed that GCK deficiency markedly reduced the expression of enzymes critical for fatty acid and cholesterol biosynthesis (Figures 5D and S4B). Conversely, GCK overexpression in PC3 cells significantly upregulated these genes (Figure S4C). Given that GCK is required for MYCN-driven neuroendocrine lineage reprogramming and that MYCN has been reported to promote lipid production^28^, we further assessed whether GCK mediates this MYCN-associated lipid metabolic program. In MYCN-overexpressing PC3 cells, knockdown of GCK effectively abolished the MYCN-induced upregulation of lipid metabolism-related genes (Figure 5E), indicating that GCK functions on the downstream of MYCN to support lipid biosynthesis. In parallel, overexpression of GCK in adenocarcinoma cells led to a marked increase in intracellular NADPH levels and NADPH/NADP⁺ ratios (Figures 5F, 5G, S4D, and S4E), a key redox cofactor required for fatty acid and cholesterol synthesis. Since NADPH is primarily produced from glucose metabolism and provides reducing power for anabolic biosynthesis, these findings suggest that GCK promotes metabolic coupling between glycolysis and lipogenesis. Consistently, GCK-deficient LASCPC-01 cells displayed a significant reduction in intracellular NADPH levels, whereas re-introduction of shRNA-resistant GCK restored NADPH to near-control levels (Figures 5H, 5I, and S4F). To further substantiate the role of GCK in lipid metabolism, we performed Oil Red O staining, which showed that knockdown of GCK markedly decreased, whereas re-expression of shRNA-resistant GCK restored, lipid droplet accumulation in LASCPC-01 cells (Figures 5J and 5K). Similarly, GCK overexpression in PC3 cells increased lipid droplet formation, as demonstrated by both Oil Red O and BODIPY 493/503 staining (Figures S4G–J). Altogether, these findings uncover a previously unrecognized function of GCK in promoting lipid metabolism in NEPC cells, highlighting its role as a metabolic–regulatory hub that bridges glucose utilization and lipid biosynthesis.

**Figure 5.**
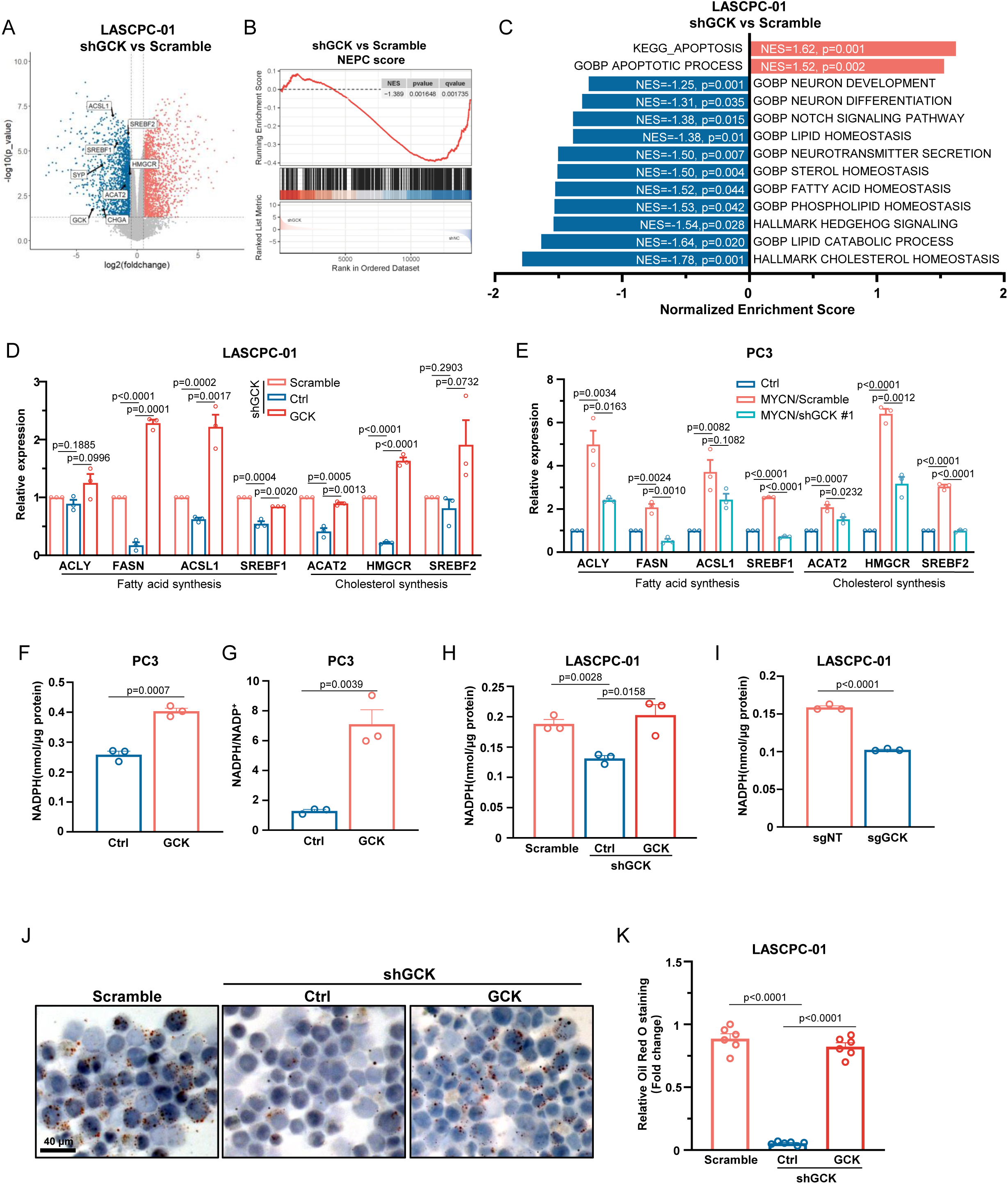
GCK promotes lipid metabolism in NEPC. (A) Volcano plot showed the differentially expressed genes in the RNA-sequencing data of LASCPC-01/scramble and LASCPC-01/shGCK. Red plots indicated significantly upregulated genes, and blue plots indicates significantly downregulated genes in LASCPC-01 shGCK versus LASCPC-01/scramble (p<0.05, FC>1.5). (B) Geneset enrichment analysis of differentially expressed genes using NEPC score geneset^27^ in LASCPC-01/shGCK versus LASCPC-01/scramble. (C) Pathway enrichment analysis of differentially expressed genes in LASCPC-01/shGCK versus LASCPC-01/scramble. (D) QPCR showing relative mRNA expression of fatty acid synthesis-related genes (*ACLY*, *FASN*, *ACSL1*, and *SREBF1*) and cholesterol synthesis-related genes (*ACAT2*, *HMGCR*, and *SREBF2*) in LASCPC-01/scramble, LASCPC-01/shGCK, and LASCPC-01/shGCK/GCK (n = 3 biologically independent experiments). (E) QPCR showing relative mRNA expression of indicated genes in PC3/Ctrl, PC3/MYCN, and PC3/MYCN/shGCK #1 (n = 3 biologically independent experiments). (F) Total NADPH levels in PC3/Ctrl and PC3/GCK (n = 3 biologically independent experiments). (G) NADPH/NADP^+^ ratio levels in PC3/Ctrl and PC3/GCK. (H) Total NADPH levels in LASCPC-01/scramble, LASCPC-01/shGCK, and LASCPC-01/shGCK/GCK (n = 3 biologically independent experiments). (I) Total NADPH levels in LASCPC-01/sgNT and LASCPC-01/sgGCK (n = 3 biologically independent experiments). (J) Representative images in LASCPC-01/scramble, LASCPC-01/shGCK, and LASCPC-01/shGCK/GCK stained by Oil Red O. Scale bar, 40 μm. (K) Quantification of Oil Red O intensity in LASCPC-01/scramble, LASCPC-01/shGCK, and LASCPC-01/shGCK/GCK (n = 3 biologically independent experiments). Data presented as mean ± s.e.m (D, E, F, G, H, I, K). Statistical significance was determined by two-tailed unpaired Student’s test (F, G, H, I, K) or one-way analysis of variance (ANOVA) with Dunnett’s multiple comparisons (D, E).

### AKT1 directly binds to and phosphorylates GCK at S373

Since GCK’s critical function in NEPC is independent of its glucose kinase activity, we next examined its potential binding partners to uncover regulatory proteins that may influence its function. It is generally accepted that GCK activity is regulated by glucokinase regulatory protein (GCKR), which binds to GCK and facilitates its nuclear sequestration under low-glucose conditions, particularly in hepatocytes^29,30^. To determine whether this regulatory mechanism also operates in NEPC, we examined GCKR expression in our models. Interestingly, although *Gckr* was relatively abundant in the liver of NEPC model mice, it was barely detectable in tumor tissues (Figure S5A). Consistently, *GCKR* expression was minimal in NEPC cell lines compared with adenocarcinoma lines (Figure S5B). These findings suggest that GCKR is not the primary regulator of GCK in NEPC. To identify alternative regulators or interacting partners of GCK, we performed immunoprecipitation followed by mass spectrometry (IP-MS) in LASCPC-01 cells. We identified 142 GCK-interacting proteins (Table S2), among which AKT1 drew particular attention due to its well-established roles in metabolic regulation and neuroendocrine differentiation. This finding was consistent with previous studies reporting that AKT1 can directly phosphorylate and regulate HK2, another member of the hexokinase family^31^.

To test whether AKT1 also interacts with GCK, we first conducted co-immunoprecipitation (Co-IP) assays in HEK293T cells transiently co-expressing GCK-V5 and AKT1-HA, which revealed a clear interaction between the two proteins (Figure 6A). Co-IP assays in LASCPC-01 and GCK-overexpressing PC3 cells similarly demonstrated endogenous interaction between AKT1 and GCK (Figures 6B and S5C). This direct interaction was further validated by GST pull-down assays, showing that recombinant AKT1 bound to full-length GCK and its C-terminal domain (residues 171–465), but not the N-terminal region (residues 1–170) (Figure 6C). Together, these results confirm that AKT1 directly binds to GCK in NEPC cells.

**Figure 6.**
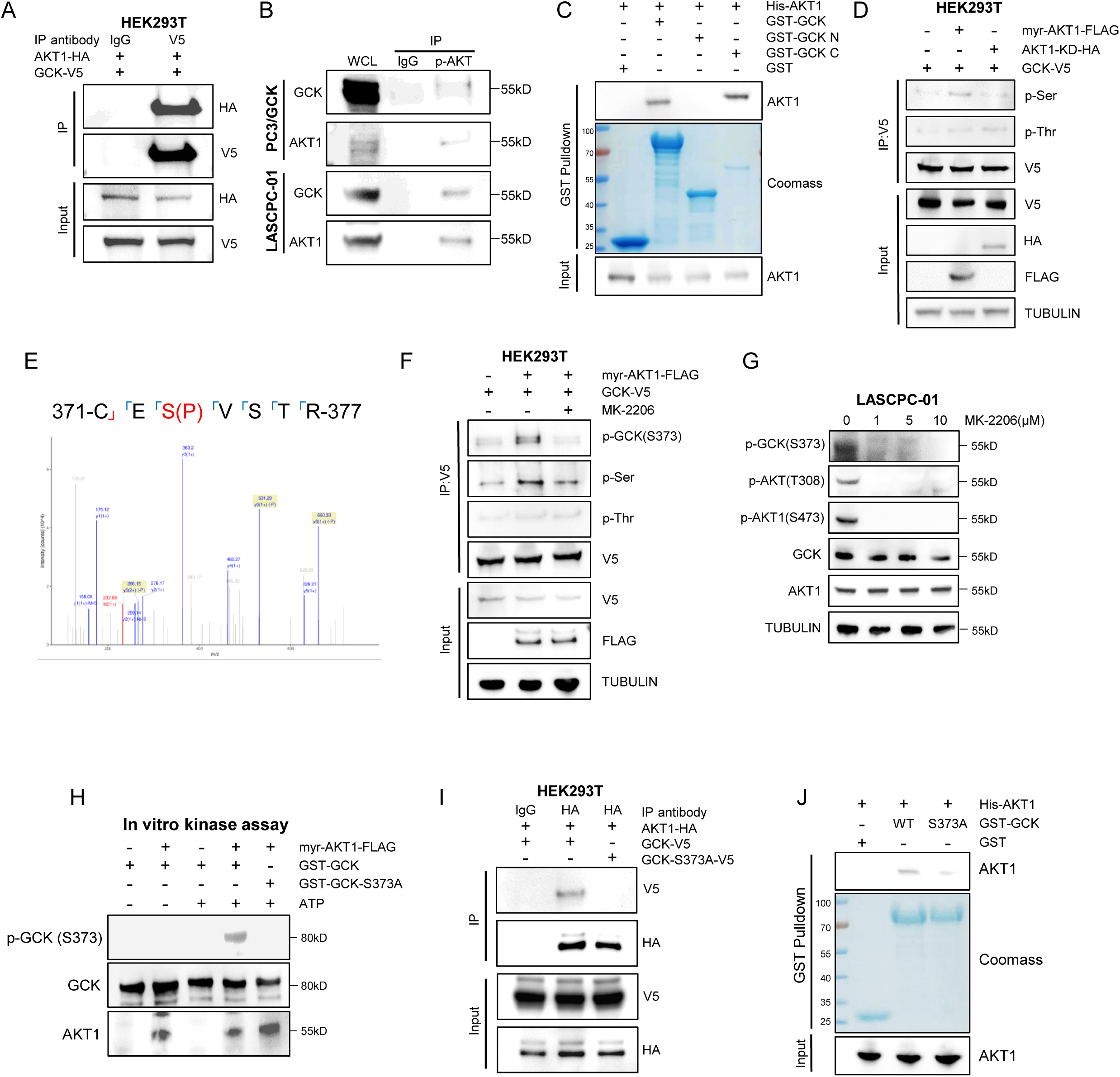
AKT1 directly binds to and phosphorylates GCK at S373. (A) Immunoprecipitation and immunoblotting assays in HEK293T cells transfected with AKT1-HA and GCK-V5 for 48 hours (n = 3 biologically independent experiments). (B) Immunoprecipitation and immunoblotting assays in PC3/GCK cells (left) and LASCPC-01 cells (right) (n = 3 biologically independent experiments). (C) GST pulldown and immunoblotting assays using bacterially purified His-AKT1 and the indicated GST-GCK (n = 3 biologically independent experiments). (D) Immunoprecipitation and immunoblotting assays in HEK293T cell with indicated transfection for 48 hours (n = 3 biologically independent experiments). (E) In vitro kinase assays followed by mass spectrometric analysis showing phosphorylation of GCK at S373. A tryptic fragment at m/z 266.15 Da (−0.93 ppm), which was matched with the +2 charged peptide 371-CES(P)VSTR-377. (F) Immunoprecipitation and immunoblotting assays in HEK293T cell with indicated transfection for 48 hours, treated with MK-2206 (1 μM) or DMSO for 12 hours (n = 3 biologically independent experiments). (G) Immunoblotting assay in LASCPC-01 cells treated with the indicated concentrations of MK-2206 for 12 hours (n = 3 biologically independent experiments). (H) In vitro kinase and immunoblotting assays of myr-AKT1-FLAG incubated with GST-GCK or GST-GCK-S373A in the presence or absence of ATP (1mM) (n = 3 biologically independent experiments). (I) Immunoprecipitation and immunoblotting assays in HEK293T cells with indicated transfection for 48 hours (n = 3 biologically independent experiments). (J) GST pulldown and immunoblotting assays using bacterially purified His-AKT1 with GST-GCK or GST-GCK-S373A (n = 3 biologically independent experiments).

Given that AKT1 is a serine/threonine kinase, we next investigated whether it phosphorylates GCK. HEK293T cells were co-transfected with GCK-V5 and either constitutively active AKT1 (myr-AKT1) or a kinase-dead mutant (K179M, AKT1-KD). Immunoprecipitation and immunoblot analyses revealed that serine residues of GCK were phosphorylated by active AKT1, whereas threonine residues showed no significant difference in phosphorylation (Figure 6D). To identify the exact phosphorylation site, we analyzed the GCK amino acid sequence using PhosphoSitePlus and iGPS, both of which predicted serine 373 (S373) as the most probable AKT1 target site (Figure S5D). This residue is evolutionarily conserved across species but distinct from corresponding positions in other hexokinase family members (Figures S5E and S5F). Consistent with these predictions, an in vitro phosphorylation assay coupled with LC-MS/MS confirmed that AKT1 phosphorylates GCK at S373 (Figure 6E). We subsequently generated a phospho-specific antibody against GCK (pS373) and verified its specificity through peptide competition assays using recombinant GCK protein (Figure S5G). Immunoprecipitation of GCK-V5 from HEK293T cells showed that co-expression with active AKT1 markedly enhanced GCK S373 phosphorylation, as detected by both phospho-GCK and pan-serine antibodies (Figure 6F). This phosphorylation was effectively blocked by MK-2206, a selective AKT inhibitor, which also reduced endogenous GCK phosphorylation in LASCPC-01 and NCI-H660 cells (Figures 6G and S5H). An in vitro kinase assay using recombinant proteins further demonstrated that wildtype GCK, but not the S373A mutant, was phosphorylated by AKT1 in the presence of ATP (Figure 6H). Moreover, Co-IP and GST pull-down assays revealed that GCK with S373A mutation lost the ability to interact with AKT1, whereas wildtype GCK maintained the interaction (Figures 6I and 6J). Importantly, mutations in GCK’s glucose-binding sites did not affect its interaction with AKT1 (Figure S5I).

Collectively, these findings demonstrate that AKT1 directly binds to and phosphorylates GCK at serine 373, and that this phosphorylation is crucial for maintaining the AKT1-GCK interaction.

### GCK functions as a protein kinase and phosphorylates AKT1

We next examined whether the interaction between GCK and AKT1 contributes to the lineage commitment of prostate cancer cells. In LASCPC-01 cells, GCK knockdown or knockout led to an unexpectedly profound reduction in AKT activity, reflected by loss of phosphorylation at T308 and S473 (Figures 7A and 7B). A similar decrease was observed in NCI-H660 cells following shRNA-mediated GCK silence (Figure 7C). Conversely, ectopic expression of GCK in LASCPC-01 cells led to a pronounced increase in AKT1 phosphorylation, particularly at S473 (Figure S6A). Consistent with these findings, overexpression of wildtype GCK, but not the S373A mutant, enhanced AKT1 phosphorylation at both T308 and S473 in response to insulin stimulation (Figure S6B).

**Figure 7.**
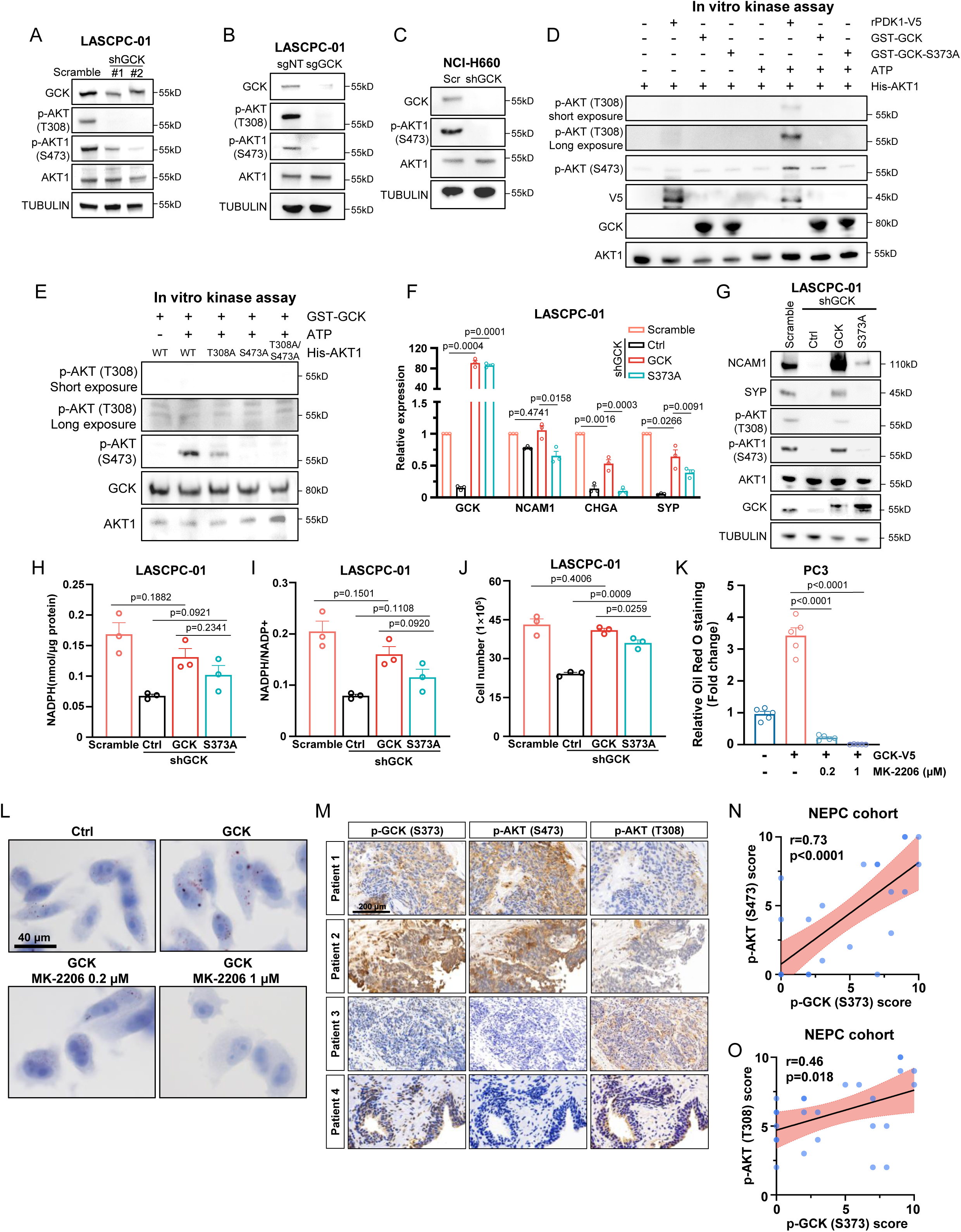
GCK functions as a protein kinase and phosphorylates AKT1. (A) Western blot showing AKT1, p-AKT(T308), and p-AKT1(S473) protein expression in LASCPC-01/scramble, LASCPC-01/shGCK #1, and LASCPC-01/shGCK #2 (n = 3 biologically independent experiments). (B) Western blot showing AKT1, p-AKT(T308), and p-AKT1(S473) protein expression in LASCPC-01/sgNT and LASCPC-01/sgGCK (n = 3 biologically independent experiments). (C) Western blot showing AKT1 and p-AKT1(S473) protein expression in NCI-H660/scramble and NCI-H660/shGCK (n = 3 biologically independent experiments). (D) In vitro kinase and immunoblotting assays of His-AKT1 incubated with indicated proteins in the presence or absence of ATP (0.5mM) (n = 3 biologically independent experiments). (E) In vitro kinase and immunoblotting assays of GST-GCK incubated with indicated His-AKT1 in the presence or absence of ATP (0.5mM) (n = 3 biologically independent experiments). (F) QPCR showing relative mRNA expression of GCK and neuroendocrine-related genes (*CHGA*, *NCAM1*, and *SYP*) in indicated LASCPC-01 cells (n = 3 biologically independent experiments). (G) Western blot showing protein expression of neuroendocrine markers (NCAM1 and SYP), GCK, p-AKT (T308), p-AKT1 (S473), and total AKT1 in indicated LASCPC-01 cells (n = 3 biologically independent experiments). (H) Total NADPH levels in indicated LASCPC-01 cells (n = 3 biologically independent experiments). (I) NADPH/NADP^+^ ratio levels in indicated LASCPC-01 cells. (J) Proliferation rate of indicated LASCPC-01 cells after 4 days of culture (n = 3 biologically independent experiments). (K) Quantification of Oil Red O intensity in PC3/Ctrl and PC3/GCK treated with the indicated concentrations of MK-2206 for 12 hours (n = 5 biologically independent experiments). (L) Representative images of Oil Red O staining in PC3/Ctrl and PC3/GCK treated with the indicated concentrations of MK-2206 for 12 hours. Scale bar, 40 μm. (M) Representative immunohistochemistry staining of p-GCK (S373), p-AKT (S473), and p-AKT (T308) in tumor samples from NEPC patients (n = 26). Representative images from four patients were shown. Scale bar, 200 μm. (N) The Pearson correlation analysis between p-AKT (S473) and p-GCK (S373) staining scores in tumor samples from NEPC patients (n = 26). (O) The Pearson correlation analysis between p-AKT (T308) and p-GCK (S373) staining scores in tumor samples from NEPC patients (n = 26). Data presented as mean ± s.e.m (F, H, I, J, K). Statistical significance was determined by two-tailed unpaired Student’s test (H, I, J, K), one-way analysis of variance (ANOVA) with Dunnett’s multiple comparisons (F), or Pearson correlation test (N, O).

Given recent evidence that several metabolic enzymes can exert “moonlighting” protein kinase activities^32,33^, these observations prompted us to hypothesize that GCK may directly phosphorylate AKT1, thereby explaining its role in lineage maintenance, NADPH generation, and lipid metabolism. To test this hypothesis, we performed an in vitro kinase assay using purified GST-GCK and His-AKT1. As shown in Figure 7D, GCK directly phosphorylated AKT1 at S473, but not at T308, the latter being the canonical site targeted by PDK1. This phosphorylation was greatly reduced when either GCK S373 or AKT1 S473 was mutated to alanine (Figures 7D and 7E), indicating that the interaction between these residues is essential for the phosphorylation event.

Importantly, the GCK S373A mutant retained normal glucose uptake and glucokinase activity, suggesting that its interaction with AKT1 is independent of GCK’s canonical enzymatic function (Figures S6C and S6D). However, unlike wildtype GCK, the S373A mutant failed to rescue neuroendocrine features in LASCPC-01 cells with endogenous GCK knockdown, nor did it promote NADPH production or lipid metabolism (Figures 7F, 7G, 7H, 7I, 7J, and S6E). Consistently, pharmacological inhibition of AKT using MK-2206 effectively abolished the GCK-induced increase in lipid metabolism, as evidenced by reduced NADPH production and diminished Oil Red O and BODIPY 493/503 staining (Figures 7K, 7L, S6F, S6H, and S6I). Furthermore, immunohistochemical analysis of NEPC tumor samples revealed a strong positive correlation between GCK S373 phosphorylation and AKT1 S473 phosphorylation (Figures 7M–7O). Together, these findings demonstrate that GCK acts as a protein kinase that directly phosphorylates AKT1 at S473, and that this GCK-AKT1 signaling axis is critical for sustaining neuroendocrine lineage identity, NADPH production, and lipid metabolism in NEPC cells.

### Targeting phosphorylation of GCK S373 via a peptidic inhibitor blocks lineage reprogramming

To explore the potential of targeting GCK S373 phosphorylation, we first analyzed the sequence surrounding the phosphorylation site and synthesized five short peptides, each fused with a cell-permeable peptide for efficient delivery^34,35^ (Figure 8A). All five peptides successfully inhibited GCK phosphorylation in LASCPC-01 cells, among which peptides 2#–5# showed complete inhibition (Figure 8B). In an in vitro kinase assay, peptide 2# and peptide #5, which contained sequences (RRACE<u>S</u>V) similar to the AKT consensus phosphorylation motif, exhibited potent inhibition of AKT1 phosphorylation (Figure 8C). Notably, peptide 2#, which demonstrated the most effective inhibition of AKT1 phosphorylation, also significantly reduced the neuroendocrine phenotype in LASCPC-01 cells (Figure 8C and 8D). Given the promising effects of peptide 2#, we proceeded to test whether it could disrupt the interaction between GCK and AKT1. Using a GST pull-down assay, we observed that peptide 2# inhibited the binding of His-AKT1 to GST-GCK in a dose-dependent manner (Figure 8E). As anticipated, peptide 2# also mitigated the phosphorylation of AKT1 at both S473 and T308 in prostate cancer cells (Figures 8F, 8G, 8H, 8I, and S7A). Interestingly, despite its effects on AKT1 phosphorylation, peptide 2# did not reduce, but rather slightly increased, glucose uptake in LASCPC-01 cells, a pattern consistent with the effects observed with GCK S373A mutant overexpression (Figures S7B and S6D).

**Figure 8.**
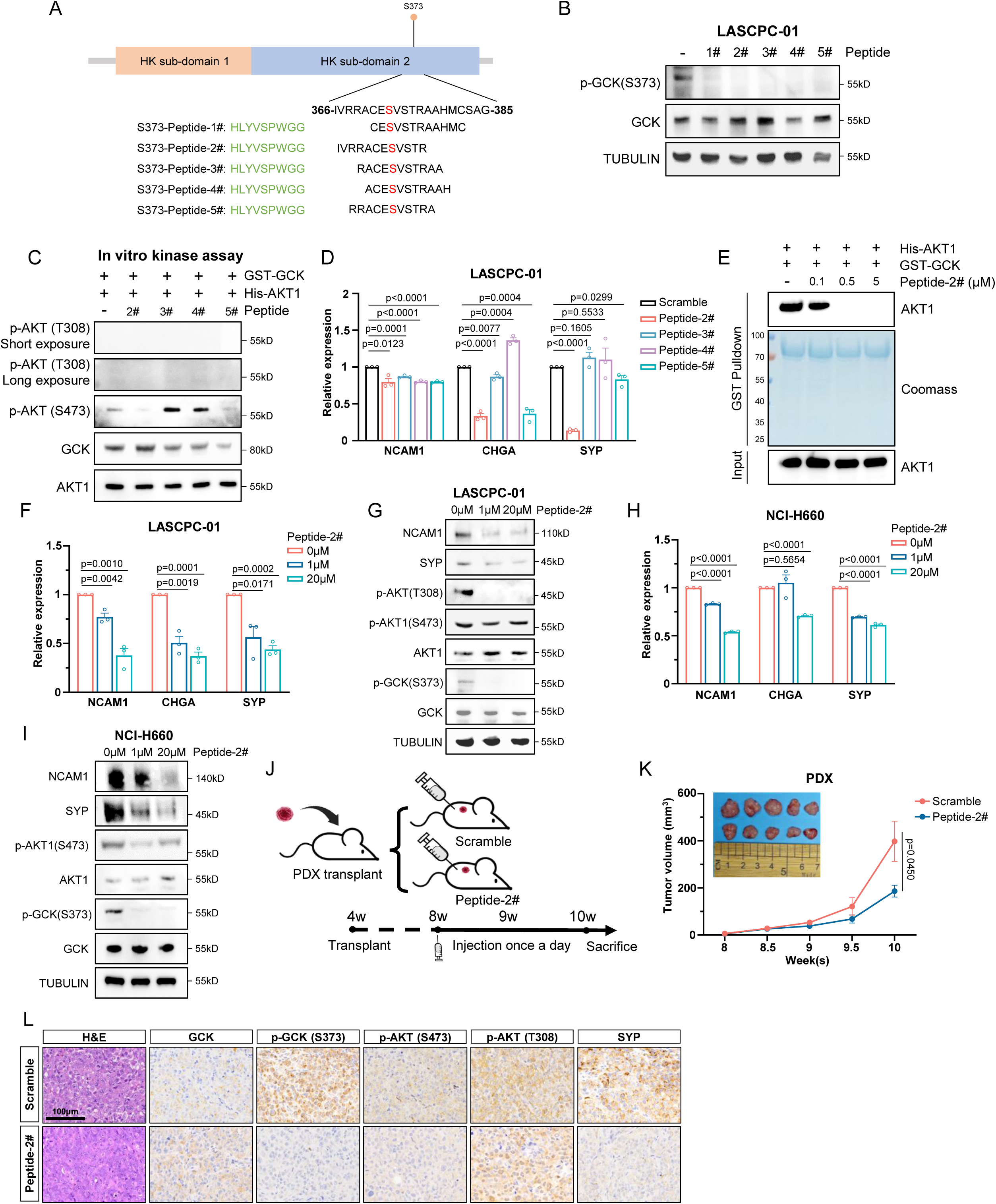
Targeting phosphorylation of GCK S373 via a peptidic inhibitor blocks lineage reprogramming. (A) Schematic illustration of designed peptides. (B) Immunoblotting analyses in LASCPC-01 cells treated with indicated peptides (n = 3 biologically independent experiments). (C) In vitro kinase and immunoblotting assays of His-AKT1 and GST-GCK incubated with indicated peptides at the presence of ATP (0.5mM) (n = 3 biologically independent experiments). (D) QPCR showing relative mRNA expression of neuroendocrine markers in LASCPC-01 cells treated with indicated peptides (n = 3 biologically independent experiments). (E) GST pulldown and immunoblotting assays using bacterially purified His-AKT1 and GST-GCK treated with the indicated concentrations of peptide 2# (n = 3 biologically independent experiments). (F) QPCR showing relative mRNA expression of neuroendocrine markers in LASCPC-01 cells treated with the indicated concentrations of peptide 2# (n = 3 biologically independent experiments). (G) Immunoblotting analyses in LASCPC-01 cells treated with the indicated concentrations of peptide 2# (n = 3 biologically independent experiments). (H) QPCR showing relative mRNA expression of neuroendocrine markers in NCI-H660 cells treated with the indicated concentrations of peptide 2# (n = 3 biologically independent experiments). (I) Immunoblotting analyses in NCI-H660 cells treated with the indicated concentrations of peptide 2# (n = 3 biologically independent experiments). (J) Schematic illustration of nude mice subcutaneously transplanted with NEPC PDX under the treatment of scramble or peptide 2#. (K) Tumor volumes of nude mice subcutaneously transplanted with NEPC PDX under the treatment of scramble or peptide 2#. Tumor volume was measured twice a week at indicated time points (n = 5 per group). (L) Representative H&E and immunohistochemistry staining of GCK, p-GCK (S373), p-AKT (S473), p-AKT (T308), and SYP in NEPC PDX treated with scramble or peptide 2#. Scale bar, 100 μm. Data presented as mean ± s.e.m (D, F, H, K). Statistical significance was determined by two-tailed unpaired Student’s test (F, H) or one-way analysis of variance (ANOVA) with Dunnett’s multiple comparisons (D, K).

Having established the potent in vitro activity of peptide 2#, we next evaluated it in vivo efficacy. Using the Pb-Cre4: *Pten*^fl/fl^; *Trp53*^fl/fl^; *Rb1*^fl/fl^ mouse model and NEPC patient-derived xenografts (Figures 8J and S7C), we found that peptide 2# treatment led to partial suppression of tumor growth and prolonged survival, without evident liver toxicity (Figures 8K, S7D, S7E, S7F, and S7G). Immunohistochemical analysis of tumor tissues revealed that peptide 2# treatment effectively reduced GCK S373 and AKT1 S473 phosphorylation, while also decreasing neuroendocrine lineage features (Figures 8L and S7H).

Taken together, these findings demonstrate that targeting the interaction between GCK and AKT1 by inhibiting the phosphorylation of GCK at S373 can block lineage reprogramming and inhibit the progression of NEPC.

## Discussion

Tumor cells accelerate glucose metabolism to meet the high energy demands of rapid proliferation and progression. Although the role of the hexokinase family in glucose import into tumor cells is well established, how these enzymes interplay with oncogenic pathways and operate a tightly regulated network between metabolic activities and malignant behaviors still remain unclear. Herein, we investigate the mechanisms underlying high glucose uptake in NEPC and demonstrate that GCK is upregulated by MYCN transcriptional regulation. Notably, we reveal that GCK interacts with AKT1 in a reciprocal regulatory loop, in which AKT1 phosphorylates GCK at S373, and GCK in turn phosphorylates and activates AKT1. To further explore this interaction, we designed a cell-penetrating peptide that targets the phosphorylation of GCK S373 and thereby disrupts the GCK-AKT1 interaction. This peptide moderately yet significantly suppressed NEPC tumor growth while remarkably blocking the lineage switch both in vitro and in vivo (Figures 8F and 8K). In contrast, complete deletion of GCK exerted a much more dramatic inhibitory effect on tumor growth (Figures 2A and 2K). Given that MYCN together with sustained AKT1 activation can transform normal prostate epithelium into NEPC^18^, and that MYCN overexpression alone is sufficient to convert prostate adenocarcinoma cells into NEPC^23^, our findings position GCK as a critical mediator interfacing MYCN and the hyperactive AKT1. Moreover, our work highlights the dual functions of GCK in NEPC progression: its metabolic role facilitates glucose uptake and provides the energy required for rapid cell proliferation, while its non-metabolic role maintains lineage commitment via a positive feedback loop with AKT1.

Such multifaceted functions of glucose metabolic enzymes have garnered considerable attention in cancer research^36^. A recent study suggests that nuclear-translocated GCK can function as a protein kinase, phosphorylating TAZ to promote tumor growth^37^. Similarly, other studies have shown that HK2 can modulate various tumor cell activities through interactions with proteins like IκBα, CD133, GSK3, and mTORC1^38–41^. These glucose-metabolizing enzymes are finely tuned in response to both intracellular and extracellular signaling cues. In the liver, GCK activity is tightly regulated by transcriptional networks such as FOXO1 in response to insulin signaling^42^, while at the protein level, high glucose promotes dissociation of the nuclear GCK–GCKR complex and translocation of GCK to the cytoplasm, where its activity is further enhanced through interaction with BAD protein located on the mitochondria^43^. Our studies, along with others, have reported multiple post-translational modifications of HK2, including deSUMOylation by SENP1^44^, phosphorylation by AKT^45^, and ubiquitination by HectH9^46^. The multifunctionality of these enzymes and their coordination with key oncogenic pathways suggest an evolutionarily advantageous strategy that tumor cells adapt to coordinate metabolic control with essential oncogenic signaling, facilitating aggressive tumor expansion under metabolic constraints. In our study, the upregulation of GCK endows NEPC cells with a robust ability to take up glucose, alongside maintaining neuroendocrine lineage through mutual reinforcement with AKT1.

Another key observation from our study was that GCK may promote lipid metabolism in NEPC cells, as evidenced by increased lipid staining and the transcriptional upregulation of lipid-related enzymes (Figures 5D and 5J). These findings align with previous reports showing that GCK activators can lead to adverse effects, such as hyperlipidemia and hepatic fat accumulation^47,48^. Conversely, inactivation of GCK has been shown to alleviate lipid accumulation in hepatocytes^49^. While our study focuses on the role of GCK in NEPC cells, we do not extend these functions and mechanisms to other endocrine or endocrine-like cells, such as pancreatic β cells and hepatocytes, where GCK is also highly expressed. Although we provide evidence for the reciprocal regulation between GCK and AKT1, we cannot exclude the possibility that GCK may also interact with other key lipid-related factors, contributing to lipid homeostasis. Given the critical role of GCK in glucose metabolism and its function in various endocrine cells, we are optimistic that the implications of our study extend beyond NEPC and may be relevant to metabolic diseases. Further investigation is needed to characterize the regulation and functions of GCK in both its metabolic and non-metabolic roles.

In conclusion, our study uncovers a previously unrecognized mechanism through which metabolic adaptation is intertwined with lineage identity in NEPC. We demonstrate that NEPC exhibits elevated glucose uptake and identify GCK as a MYCN-induced metabolic enzyme indispensable for sustaining the neuroendocrine lineage. Beyond its canonical role, GCK promotes lipid metabolism and functions as a protein kinase, forming a reciprocal phosphorylation loop with AKT1—AKT1 phosphorylates GCK at S373, whereas GCK phosphorylates AKT1 at S473. Together, these findings delineate a noncanonical kinase activity of GCK and establish the AKT1-GCK axis as a pivotal regulator linking metabolic reprogramming and neuroendocrine lineage plasticity in prostate cancer.

## Experimental model and subject details

### Ethics

The research complied with all relevant ethical regulations. The collection of human samples and research conducted in this study was approved by the Research Ethics Committee of the Renji Hospital, Shanghai Jiao Tong University School of Medicine (approval numbers: RA-2024-081). Clinical samples and information were collected after written informed consent. All animal experiments were performed in compliance with the Guide for the Care and Use of Laboratory Animals (National Academies Press, 2011) and were approved by the Ren Ji Hospital Laboratory Animal Use and Care Committee.

### Cell culture

The human prostate cancer cell lines LNCaP (SCSP-5021), PC3 (SCSP-532), and DU145 (SCSP-5024) cells were purchased from the Cell Bank, Shanghai Institutes for Biological Sciences, Chinese Academy of Sciences, cultured in RPMI-1640 medium or DMEM (Gibco) supplemented with 10% fetal bovine serum (Invitrogen), 100 U/mL penicillin, and 100 μg/mL streptomycin (Gibco). LASCPC-01 (CRL-3356) and NCI-H660 (CRL-5813) cells were purchased from ATCC, cultured in HITES medium, supplemented with 5% fetal bovine serum (Invitrogen), 100 U/mL penicillin, and 100 μg/mL streptomycin. All the cells were incubated at 37°C in a humidified incubator with 5% CO_2_ and routinely tested for mycoplasma. The identity of the cell lines was verified through high-resolution small tandem repeats (STR) profiling.

### Animals

All animal experiments and procedures were carried out in strict accordance with the Guidelines for the Care and Use of Laboratory Animals set by the U.S. National Institutes of Health (National Academies Press; 2011) and were performed following the ethical guideline approved by Ethics Committee of Ren Ji Hospital, School of Medicine, Shanghai Jiao Tong University.

C57BL/6 background Pb-Cre4: *Pten*^fl/fl^; *Trp53*^fl/fl^ mice were generated by breeding; Pb-Cre4^−^: *Pten*^fl/fl^; *Trp53*^fl/fl^ females with Pb-Cre4^+^: *Pten*^wt/fl^; *Trp53*^fl/fl^ males. C57BL/6 background Pb-Cre4: *Pten*^fl/fl^; *Trp53*^fl/fl^; *Rb1*^fl/fl^ mice were obtained from crossing Pb-Cre4^-^: *Pten*^fl/fl^; *Trp53*^fl/fl^; *Rb1*^fl/fl^ females with Pb-Cre4^+^: *Pten*^wt/fl^; *Trp53*^fl/fl^; *Rb1*^fl/fl^ males. All mice were born and maintained under pathogen-free conditions. All genotyping was done by PCR.

For the xenograft experiments, male BALB/c nude mice were provided by the animal laboratory of Ren Ji Hospital. To evaluate the influence of GCK in NEPC, we established LASCPC-01 xenograft models. In brief, LASCPC-01/scramble and LASCPC-01/shGCK cells (5 x 10^6^ cells in 100 μL DMEM/Matrigel) were implanted into the right armpit of male BALB/c nude mice. Three weeks after inoculation, the tumor volume of xenografts was measured every 3 days. The mice were sacrificed at week 8 and tumors were collected and processed for downstream analyses by overnight fixation with 10% buffered formalin followed by tissue processing and embedding in paraffin.

### NEPC patient-derived xenograft model

NEPC patient-derived xenografts (PDX) were subcutaneously transplanted into 4-week-old male BALB/c nude mice. Mice bearing tumors of 50-100mm^3^ were randomly assigned into two groups: Scramble group and GCK S373 peptide group (5 mg.kg-1). The treated mice were intraperitoneally injected once a day. Tumor volume (calculated as Volume = 0.52 x Length x Width^2^) was measured using an electronic vernier caliper every 3 days. The mice were sacrificed after three weeks of treatment and tumors were collected and processed for downstream analyses by overnight fixation with 10% buffered formalin followed by tissue processing and embedding in paraffin.

### NEPC patient-derived organoid model

The generation of NEPC patient-derived organoids (PDO) derived from NEPC PDX tumor. Tumor were harvested, minced, and digested in DMEM containing 2% FBS, 0.2 mg/mL collagenase Ⅳ, 0.2 mg/mL collagenase I, 0.1 mg/mL dispase, and 0.01 mg/mL Dnase I at 37 °C for 2 h. Organoid were cultured in William’s E medium supplemented with 5% Matrigel, 5% charcoal-stripped FBS, 10 μM Y-27632, 10 ng/mL epidermal growth factor, 100 nM DHT and 1× Glutamax in a low adsorption 96-well plate.

### Method details

#### 18F-FDG PET Imaging in vivo

In vivo small animal imaging was conducted at Nuclear Medicine Department of Renji Hospital, Shanghai Jiaotong University School of Medicine. Mice were fasted for 8 hours and injected with approximately 100 μCi of ^18^F-FDG via lateral tail vein (the exact dose was calculated by measuring the syringe before and after injection). Mice were anaesthetized with isoflurane for an hour. Then, mice were imaged by micro-PET/CT (Inviscan). ^18^F-FDG uptake was quantified by drawing region of interest (ROI) using IRIS PET/CT software and plotting maximum uptake values (SUVmax).

### Western blot

Cells and tissues were lysed in 1% SDS lysis buffer (Beyotime), and the protein concentration was determined by BCA Protein Assay (Thermo Fisher Scientific). Equal amounts of protein were separated by 10% SDS-PAGE gel and transferred to 0.45 μm PVDF membranes (Millipore). The membranes were blocked with 5% BSA in TBST and incubated with specific antibodies at 4°C overnight. Appropriate secondary antibodies were then used, and an ECL detection system (Tanon, China) was used to detect the protein bands.

### Quantitative PCR

Total RNA from cells and tissues were extracted with TRIzol Reagent (15596026, Invitrogen), and the concentration of RNA was quantified by a NANODROP 2000c spectrophotometer (Thermo Fisher Scientific). The reverse transcription was carried out by Reverse transcription kit (R223-1, Vazyme, China). The qPCR was performed using SYBR-green (Q711-2, Vazyme, China) in a LightCycler 480 qPCR machine (Roche).

Relative gene mRNA expression level was determined using the 2^-ΔΔCq^ method. The primer sequences used to amplify the target genes are listed in Table S3.

### Glucokinase activity assays

Glucokinase activity was measured by monitoring the rate of NADH formation using a G6PDH-coupled reaction as previously described^50,51^. Briefly, GCK-V5 purified from HEK293T cells was incubated with reaction buffer (100 mM HEPES pH 7.4, 150 mM KCl, 6 mM MgCl2, 1 mM DTT, 1 mM NAD, 0.05% BSA, 2.5 units G6PDH) and performed at 37 °C in 100 μL total volume per well for 30 min. Absorbance at 340 nm was recorded every five min between 30 and 60 minutes. To determine glucose-dependent kinetic parameters, glucose was varied (0–100 mM) while maintaining ATP at 5 mM.

### In vitro kinase assays

In vitro kinase assays were performed as reported^52^. In brief, for the AKT in vitro kinase assay, purified active myr-AKT-FLAG from HEK293T was incubated with bacterially purified GST-GCK or GST-GCK-S373A (200 ng) in 25 μL kinase buffer (25 mM Tris-HCl (pH 7.5), 5 mM beta-glycerophosphate, 2 mM dithiothreitol (DTT), 0.1 mM Na3VO4, 10 mM MgCl2, and 1mM ATP) at 30 °C for 1 hour. For GCK in vitro kinase assay, bacterially purified GST-GCK, purified GCK-V5 or purified PDK1-V5 from HEK293T was incubated with bacterially purified His-AKT1 and other mutations in the presence of ATP for 1 hour. The reaction was terminated by adding SDS–PAGE loading buffer and heating to 100 °C for 5 minutes. The reaction mixture was then subjected to SDS–PAGE analysis.

### Glucose uptake assay

Glucose uptake of cells was measured by Glucose Assay Kit-WST (Dojindo) according to the manufacturer’s instructions. In brief, 5 x 10^5^ cells were seeded in each well of 6-well plate. After incubated for 36 hours, culture medium was collected and incubated with reaction buffers for 30 minutes at 37 °C. The absorbance was measured at 450 nm. Glucose uptake was normalized to the average protein concentration of cells before the incubation.

#### NADP+/NADPH Assay

NADPH concentration of cells was measured by NADP+/NADPH assay kit (Beyotime Biotechnology) according to manufacturer’s instructions. If brief, 5 x 10^5^ cells were collected and lysed. The supernatant was heated at 60 °C for 30 minutes to decompose NADP+. Then, the supernatant was incubated with reaction buffer for 10 minutes at 37 °C to convert NADP+ to NADPH. The absorbance was measured at 450 nm. The amount of NADPH was normalized to the average protein concentration of cells before the incubation.

### GST pulldown assay

Equal amounts of His-tagged purified protein (200 ng per sample) were incubated with 100 ng of GST fusion proteins together with glutathione agarose beads in a modified binding buffer (20 mM pH7.4 Tris-HCl, 150 mM NaCl, 1 mM EDTA, 0.5% NP-40, 1 mM PMSF, 100 μM phenylmethylsulfonylfluoride, 100 μM leupeptin, 1 μM aprotinin, 100 μM sodium orthovanadate, 100 μM sodium pyrophosphate and 1 mM sodium fluoride). The glutathione agarose beads were then washed six times with binding buffer and then added SDS-PAGE loading buffer and heating to 100 °C for 5 minutes. The reaction mixture was then subjected to immunoblotting analysis.

### IP and immunoblotting analysis

The extraction of proteins using a modified buffer from cultured cells was followed by IP and immunoblotting using corresponding antibodies. Briefly, the cells were collected and washed with cold PBS three times. Cell pellets were resuspended and lysed in the lysis buffer (20 mM pH7.4 Tris-HCl, 150 mM NaCl, 1 mM EDTA, 0.5% NP-40, and protease inhibitor cocktail) for 30 minutes at 4 °C. The lysates were centrifuged at 15,000g, and supernatant was transferred to a pre-chilled microcentrifuge tube. Protein concentration was examined using the BCA Protein Assay Kit (Pierce) according to the instructions of the manufacturer. About 10% of the supernatant was collected for western analysis as inputs. A total of 1,500 μg of protein was incubated with indicated antibodies overnight and then mixed with protein A/G–magnetic beads for 2 hours. The IP beads were washed with lysis buffer five times, followed by immunoblotting analysis. The blots were blocked with 5% bovine serum album followed by the incubation of primary antibodies and HRP-conjugated secondary antibodies.

### Immunofluorescence analysis

Cells were fixed by 4% formaldehyde for 10 minutes, permeabilized by 0.1% Triton X-100 for 10 minutes and blocked with 5% BSA/PBS at room temperature for 1 hour. Primary antibodies were then incubated at 4 °C overnight, followed by rigorous wash with PBS. Double immunofluorescence staining was performed using the Alexa Fluor™ 555 Tyramide SuperBoost™ Kit (B40913, Thermo Fisher Scientific) and Alexa Fluor™ 488 Tyramide SuperBoost™ Kit (B40922, Thermo Fisher Scientific) according to the provided manufacturer’s instructions. The samples were then washed rigorously with PBS. DAPI (1:1000) was added for DNA staining. Images were taken using Nikon-80i microscope under 40x objective. To quantify immunofluorescence staining, two independent researchers calculated the average number of membrane-positive cells in eight to nine random 40x fields.

### Lipid droplet staining

Cells were fixed with 4% formaldehyde for 10 minutes, then washed rigorously with PBS. For BODIPY staining, cells incubated with PBS containing 1 mM BODIPY in the dark at RT for 30 minutes, then washed three times with PBS. DAPI (1:1000) was added for DNA staining. For Oil red O staining, cells were washed with 60% isopropanol for 1 minute and stained with 0.3% Oil red O at RT for 30 minutes, then washed three times with 60% isopropanol. The lipid droplet numbers per cell were calculated based on the observations of at least 5 high power field.

### Chromatin immunoprecipitation (CHIP) assay

ChIP assay was performed using an SimpleChIP Plus Enzymatic Chromatin IP Kit (9005, Cell Signaling Technology) according to the manufacturer’s instructions. Antibodies used in the ChIP assay are listed in Table S3. Briefly, cells were crosslinked with 10% formaldehyde, then sonicated to an average length of 250–1000 bp. Then the precipitated chromatins were decross-linked and amplified by qPCR. All the CHIP primer sets are listed in Table S3.

### Immunohistochemistry (IHC) analysis

Prostate tumor tissues were fixed with 10% formalin for 24 hours, then embedded in paraffin and cut into 4 μm series sections. Sections were subsequently deparaffinized, rehydrated, and boiled for antigen retrieval (EDTA, pH 8.0). Sections were blocked with 10% donkey serum, then incubated with primary antibodies overnight at 4°C. Antibodies used in the IHC analysis are listed in Key resources table. The slides were then incubated by corresponding secondary antibodies for 60 minutes at room temperature. The sections were incubated by diaminobenzidine (DAB) and counterstained with hematoxylin. Representative images were taken using a Nikon light microscope.

To detect apoptotic cells, terminal deoxynucleotidyl transferase (TdT)-mediated dUTP digoxigenin nick-end labeling (TUNEL) assay was performed with TUNEL Apoptosis Assay Kits (Beyotime) according to the manufacturer’s instructions.

For immunostaining quantification, the H-score system was obtained by multiplying staining intensity (0 for no staining, 1 for weak staining, 2 for moderate staining, and 3 for intense staining) by the percentage (0–100) of cells showing that intensity.

For antibody verification with p-GCK(S373) protein, anti-p-GCK(S373) antibody (10ng) was incubated with purified p-GCK(S373) protein (100 ng) at 4°C overnight, then followed by immunohistochemistry to validate antibody specificity.

### Cell viability Assays and Dose-response Assays

Cell Counting Kit-8 assay was carried out to assess the viability of indicated prostate cancer cells. Six thousand cells of each treated prostate cell line were seeded into 96-well plates, and 10 μL CCK-8 reagent (Vazyme) was added to each well at the appropriate time point. Then the plates were incubated in the dark incubator for 2 hours. The absorbance values at 450 nm were measured with Thermo Fisher Scientific SkanIt Software 4.1.

### Plasmids and lentiviral production

All plasmids were verified by DNA Sanger sequencing. Lentivirus was prepared using a three-plasmid packing system. Briefly, PLKO.1 or pLVX-304 vectors were co-transfected into HEK293T cells along with expression vectors containing the psPAX2 and pMD2G genes.

Lentivirus was harvested at 48 and 72 hours after transfection, and the virus was cleansed by the 0.45 μm filter. Stable cell lines were selected out in 1 μg/mL puromycin for 1 week, 10 μg/mL blasticidin for 2 weeks, or 100 μg/mL hygromycin B for 2 weeks.

To generate single-cell clones with endogenous GCK promoter-deficient cells, a CRISPR/Cas9 approach was conducted. A Cas9-stable HEK293T/sgCtrl cells were generated with lenticrispr-v2 (83480, Addgene). Then, we designed two guide RNAs (listed in Table S3) targeting 80 bp sequence in the promoter region of GCK, and transfected them into HEK293T/sgCtrl cells. Single-cell clone selection was initiated 48 hours after transduction.

### Dual-luciferase reporter assay

HEK293T cells were seeded onto 24-well plates (1 x 10^5^ cells per well) and allowed to grow overnight. Then the transient transfection of indicated plasmids containing Renilla (E1910, Promega) was carried out using Lipofectamine 3000 (Invitrogen). After 48 hours, luciferase activity was conducted with the Dual Reporter Luciferase Assay System (E1910, Promega), according to the manufacturer’s instruction. The relative levels of luciferase activity were normalized to the levels of Renilla luciferase activity and the control group 48 hours after transfection.

### Mass Spectrometry analysis

For identification of interacting proteins, a protein band visualized via Coomassie blue staining was excised from an SDS–PAGE gel and digested in gel in 50 mM ammonium bicarbonate buffer containing 1 μg trypsin overnight at 37 °C. LC-MS/MS analysis was performed on a Thermo Scientific Q Exactive HF mass spectrometer (Thermo Fisher Scientific) that was coupled to HPLC (EASY-nLC 1000 HPLC system, Thermo Fisher Scientific). Protein identification required at least one unique or razor peptide per protein group. Quantification in MaxQuant (1.6.0.1 version) was performed using the label free quantification (LFQ) algorithm. Persues software (1.6.0.2 version) was used to process relative quantification and biostatistics analysis.

### RNA sequence analysis

Total RNA was extracted using the TRIzol reagent according to the manufacturer’s protocol. RNA purity and quantification were evaluated using the NanoDrop 2000 spectrophotometer (Thermo Scientific, USA). RNA integrity was assessed using the Agilent 2100 Bioanalyzer (Agilent Technologies, Santa Clara, CA, USA). Then the libraries were constructed using TruSeq Stranded mRNA LT Sample Prep Kit (Illumina, San Diego, CA, USA) according to the manufacturer’s instructions. The transcriptome sequencing and analysis were conducted by OE Biotech Co., Ltd. (Shanghai, China). The libraries were sequenced on an Illumina HiSeq X Ten platform and 150 bp paired-end reads were generated. Differential expressed genes between the two groups were calculated in R v4.0.5 by limma package v3.46.0. P < 0.001 with log2FoldChange > 1 or < -1 was set as cutoff. Gene set enrichment analysis was performed on GSEA function of clusterProfiler package v3.18.1 or GSEA software v4.1.0 to analyze the biological difference. The enrolled gene sets included Gene Ontology, Hallmark, and Kyoto Encyclopedia of Genes and Genomes adopted from the Molecular Signatures Database.

### Single-cell RNA sequence reanalysis

The raw gene expression matrix of single-cell transcriptome was derived from GSE137829 and further analyzed in R 4.0.5 using Seurat package v4.0.2. The data from 6 castration-resistant prostate cancer samples were integrated using the canonical correlation analysis approach. Cells with all the following features were filtered after Seurat-based quality control: (1) 500–7,000 detected genes; (2) < 10% mitochondrial gene expression; (3) < 3% red blood cell gene expression. Epithelial cells were identified and extracted based on the expression level of EPCAM, KRT5, KRT8, KRT14, KRT18, and CDH1. Unsupervised cell clusters were identified by shared nearest neighbor modularity optimization-based clustering algorithm. Differentially expressed genes between the high-neuroendocrine clusters and the low-neuroendocrine clusters were calculated by using the FindMarkers function in the Seurat package. P < 0.001 with log2FoldChange >1 or <−1 was set as the cutoff.

### ALT/AST assay

Serum ALT, AST levels were tested according to the manufactures’ instructions (Nanjing Jiancheng, C009-2-1, C010-2-1, respectively). Briefly, 20 µL preheated substrate was dispensed into the detection and control wells, and 5 µL serum added to the detection wells, in triplicate. The absorbance values at 450 nm were measured with Thermo Fisher Scientific SkanIt Software 4.1.

### Peptide synthesis

All peptides were synthetized by Guoping Pharmaceutic Inc (Hefei, China). Synthetic peptides were purified to >98% purity by high pressure liquid chromatography for both in vitro and in vivo use. The amino acids of peptides used in vivo use were all D isoforms. The amino sequences are listed in Table S3.

### Statistical analysis and reproducibility

All statistical analyses were performed with GraphPad 8.0 software. Two-tailed Student’s t-test assuming equal variance was used, and one-way analysis of variance for independent variance. P < 0.05 was considered significant. Data are presented as means ± s.e.m. All animals were randomized and exposed to the same environment. Blinding was not performed in tumor measurement and IHC staining.

## Data availability

RNA-seq data have been in the GEO under the accession GSE271906. Previously published sequencing data were re-analysed here: cBioPortal database^15^, GSE90891^16^ and GSE137829^53^. The remaining data are available within the Article, Supplementary Information or Source Data file. Source data are provided with this paper.

## Abbreviations

SUVmax: Maximum standardized uptake value
NEPC: Neuroendocrine prostate cancer
CRPC: Castration resistant prostate cancer
BPH: Benign prostatic hyperplasia
DKO: Pb-Cre4: Pten^fl/fl^; Trp53^fl/fl^ mice
TKO: Pb-Cre4: Pten^fl/fl^; Trp53^fl/fl^; Rb1^fl/fl^ mice
GCK: Glucokinase
HK: Hexokinase
PDO: Patient-derived organoid
PDX: Patient-derived xenograft
BS: Binding site
GSEA: Gene set enrichment analysis
GCKR: Glucokinase regulator

## Acknowledgements

We thank Prof. Wenjing Du (School of Basic Medicine Peking Union Medical College) for the HA-tagged AKT1 kinase-dead mutant plasmid, and Prof. Jinke Cheng (Shanghai Jiao Tong University School of Medicine) for the FLAG-tagged myr-AKT1 plasmid. This work was supported by National Natural Science Foundation of China 32470785 (Q.W.), 82503953 (K.S.), Shanghai Municipal Education Commission Grant Support 2023ZKZD23 (W.X.), Renji Hospital LYZXHXKT220845 (W.X.), PNO-0106 (W.X.), and China Postdoctoral Science Foundation 2025M772209 (K.S.).

## Author contributions

Q.W. and W.X. conceived and supervised the project. K.S., R.S., and Y.J. designed and performed the core experiments, analyzed the data, and drafted the initial manuscript. W.Z., X.Liu, X.Chai, J.W., B.L., A.L., H.W., and T.W. assisted with experiments, data processing, and methodology development. X.Z., Z.J., H.H.Z., L.D., Y.Z., and B.D. provided technical support, resources, and critical feedback on experimental design and interpretation. J.P. contributed to experimental assistance and project coordination. Q.W. and W.X. revised the manuscript with input from all authors. All authors discussed the results and approved the final version of the manuscript.

---

**Figure S1.**
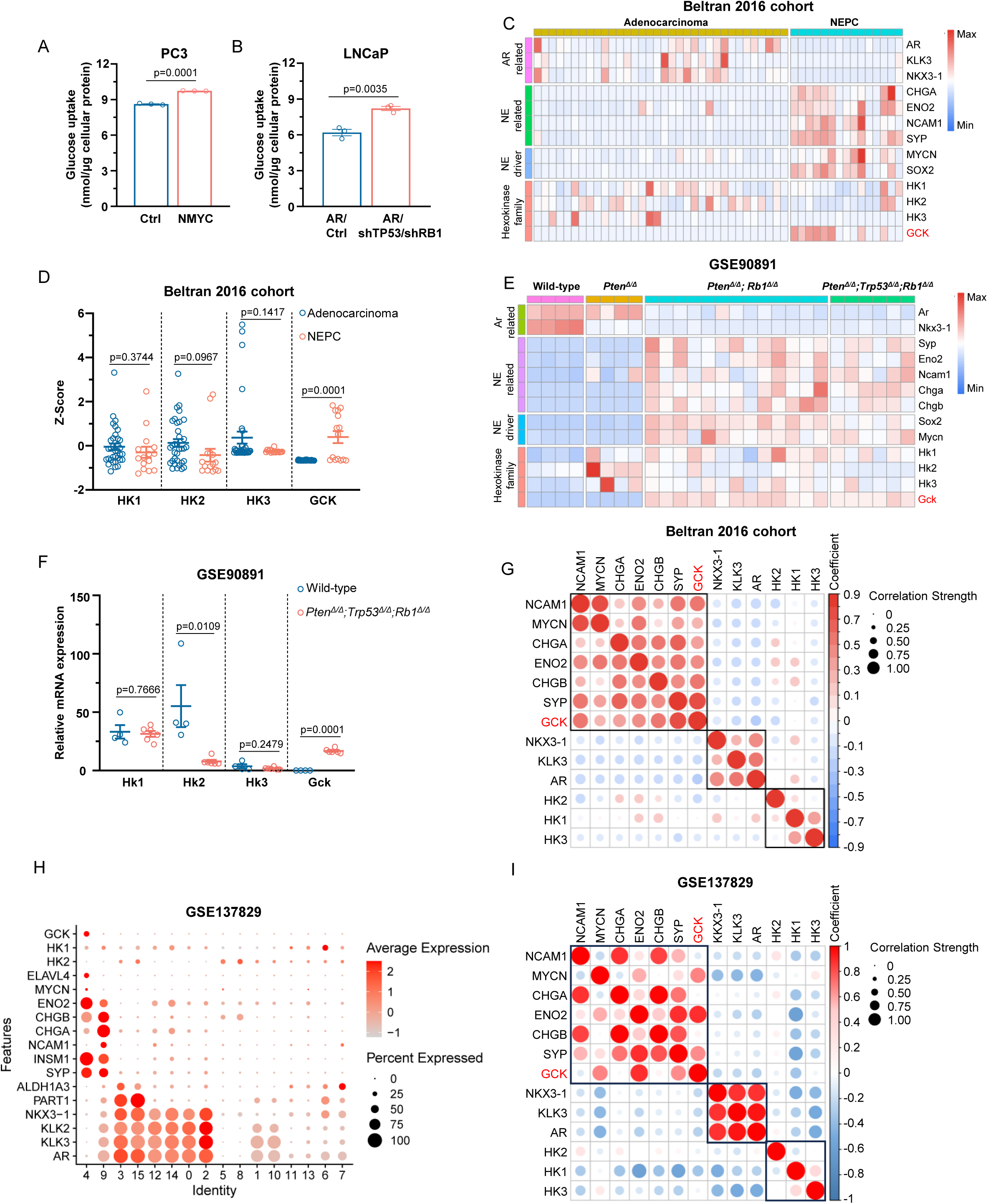
NEPC shows high glucose uptake and elevated GCK expression. (A) Glucose uptake assay in PC3/Ctrl and PC3/MYCN cells (n = 3 biologically independent experiments). (B) Glucose uptake assay in LNCaP/AR/Ctrl and LNCaP/AR/shTP53/shRB1 cells (n = 3 biologically independent experiments). (C) Heatmap showing the mRNA expression of hexokinase family, neuroendocrine-related genes, and AR-related genes in the Beltran 2016 cohort^15^ (n = 34 CRPC-adenocarcinoma samples, n = 15 CRPC-neuroendocrine samples). (D) Scatter plot showing the mRNA expression levels of the hexokinase family from CRPC-adenocarcinoma (n = 34) or neuroendocrine prostate cancer (n = 15) in the Beltran 2016 cohort. (E) Heatmap showing the mRNA expression of hexokinase family, neuroendocrine-related genes, and Ar-related genes in prostate samples from wildtype mice (n = 4), Pb-Cre4: *Pten*^fl/fl^ mice (n = 4), Pb-Cre4: *Pten*^fl/fl^; *Rb1*^fl/fl^ mice (n = 13), and Pb-Cre4: *Pten*^fl/fl^; *Trp53*^fl/fl^; *Rb1*^fl/fl^ mice (n = 6). Data obtained from GSE90891^16^. (F) Scatter plot showing the mRNA expression levels of the hexokinase family in prostate samples from Pb-Cre4: *Pten*^fl/fl^; *Trp53*^fl/fl^; *Rb1*^fl/fl^ mice (n = 6) or wildtype littermates (n = 4). Data obtained from GSE90891. (G) Pearson correlation analysis across the mRNA expression of hexokinase family, neuroendocrine-related genes, and AR-related genes. Bubble color indicates the Pearson correlation coefficient (red for positive, blue for negative), and bubble size reflects the correlation strength. Data obtained from Beltran 2016 cohort. (H) Dot plots illustrating the mRNA expression level of hexokinase family, neuroendocrine-related genes, and AR-related genes across 16 single-cell clusters from GSE137829^53^. (I) Pearson correlation analysis across the mRNA expression of hexokinase family, neuroendocrine-related genes, and AR-related genes. Bubble color indicates the Pearson correlation coefficient (red for positive, blue for negative), and bubble size reflects the correlation strength Data obtained from GSE137829. Data presented as mean ± s.e.m (A, B, D, F). Statistical significance was determined by two-tailed unpaired Student’s test (A, B, D, F).

**Figure S2.**
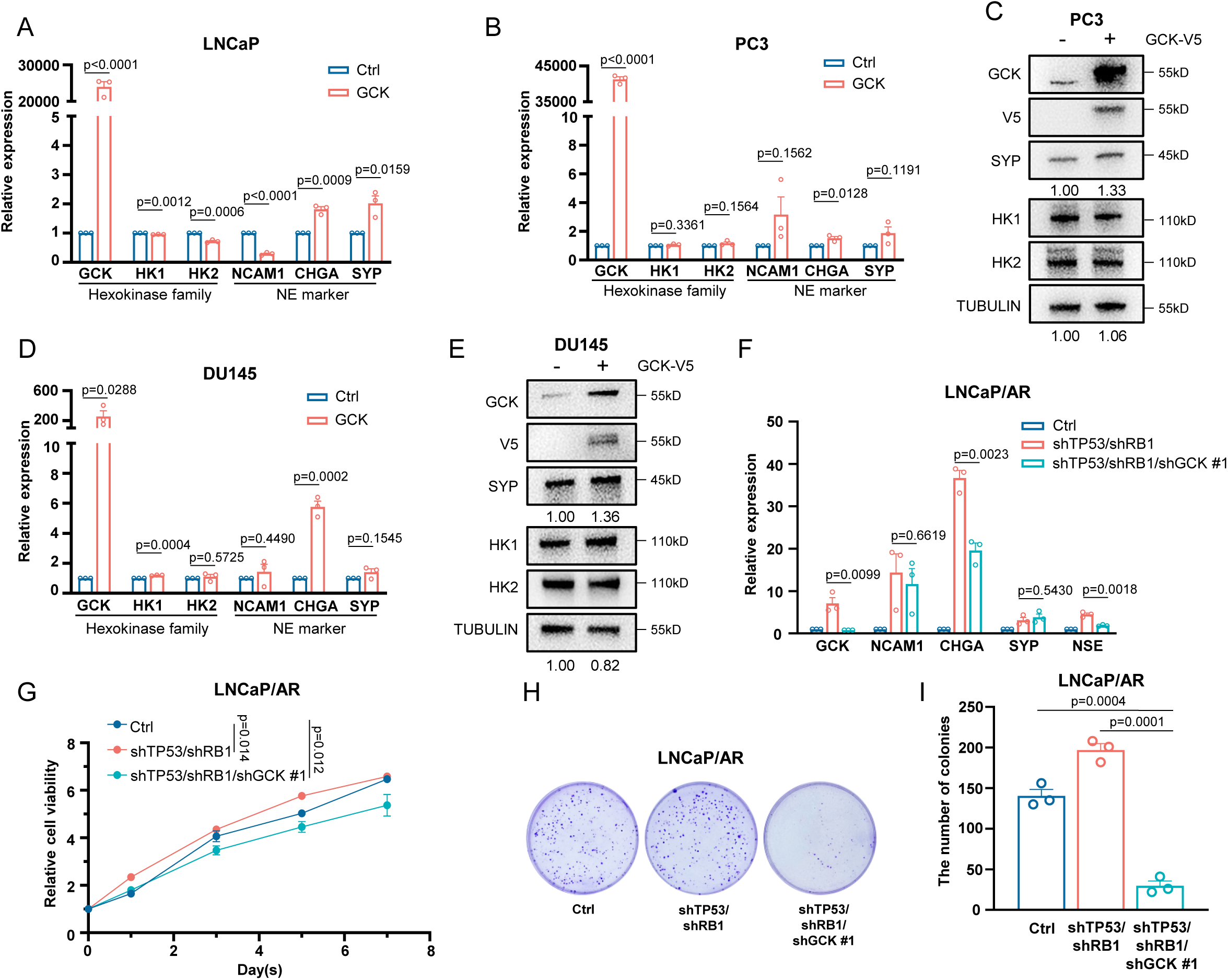
GCK is essential for maintaining the neuroendocrine lineage in prostate cancer. (A) QPCR showing relative mRNA expression of indicated genes in LNCaP/Ctrl and LNCaP/GCK (n = 3 biologically independent experiments). (B) QPCR showing relative mRNA expression of indicated genes in PC3/Ctrl and PC3/GCK. (n = 3 biologically independent experiments). (C) Western blot showing indicated protein expression in PC3/Ctrl and PC3/GCK (n = 3 biologically independent experiments). (D) QPCR showing relative mRNA expression of indicated genes in DU145/Ctrl and DU145/GCK (n = 3 biologically independent experiments). (E) Western blot showing indicated protein expression in DU145/Ctrl and DU145/GCK (n = 3 biologically independent experiments). (F) QPCR showing relative mRNA expression of indicated genes in LNCaP/AR/Ctrl, LNCaP/AR/shTP53/shRB1, and LNCaP/AR/shTP53/shRB1/shGCK #1 (n = 3 biologically independent experiments). (G) Cell viability of LNCaP/AR/Ctrl, LNCaP/AR/shTP53/shRB1, and LNCaP/AR/shTP53/shRB1/shGCK #1 at indicated time points (n = 3 biologically independent experiments). (H) Representative images of colony formation assay in LNCaP/AR/Ctrl, LNCaP/AR/shTP53/shRB1, and LNCaP/AR/shTP53/shRB1/shGCK #1. (I) Quantification of colony formation assay in LNCaP/AR/Ctrl, LNCaP/AR/shTP53/shRB1, and LNCaP/AR/shTP53/shRB1/shGCK #1 (n = 3 biologically independent experiments). Data presented as mean ± s.e.m (A, B, D, F, G, I). Statistical significance was determined by two-tailed unpaired Student’s test (A, B, D, F, I) or one-way analysis of variance (ANOVA) with Dunnett’s multiple comparisons (G).

**Figure S3.**
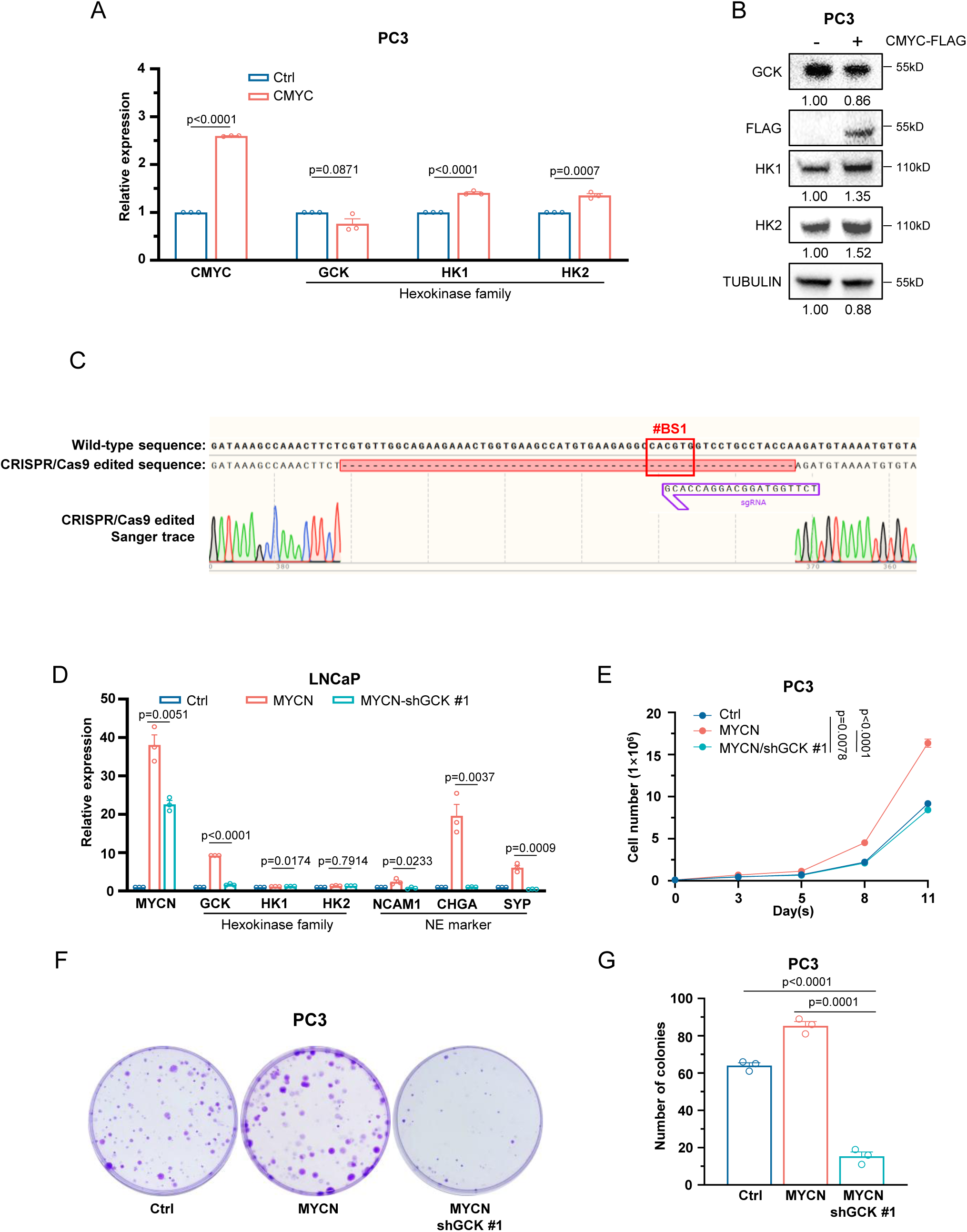
MYCN transcriptionally upregulates GCK. (A) QPCR showing relative mRNA expression of indicated genes in PC3/Ctrl and PC3/CMYC (n = 3 biologically independent experiments). (B) Western blot showing indicated protein expression PC3/Ctrl and PC3/CMYC-FLAG (n = 3 biologically independent experiments). (C) Sanger sequence of genomic DNA from PC3/sgCtrl and PC3/sgGCK promoter. The deletion mutation sequence is indicated by a solid red box. (D) QPCR showing relative mRNA expression of indicated genes in LNCaP/Ctrl, LNCaP/MYCN, and LNCaP/MYCN/shGCK #1 (n = 3 biologically independent experiments). (E) Proliferation rate of PC3/Ctrl, PC3/MYCN, and PC3/MYCN/shGCK #1 cells at indicated time points (n = 3 biologically independent experiments). (F) Representative images of colony formation assay in indicated PC3 cells. (G) Quantification of colony formation assay in indicated PC3 cells (n = 3 biologically independent experiments). Data presented as mean ± s.e.m (A, D, E, G). Statistical significance was determined by two-tailed unpaired Student’s test (A, D, G) or one-way analysis of variance (ANOVA) with Dunnett’s multiple comparisons (E).

**Figure S4.**
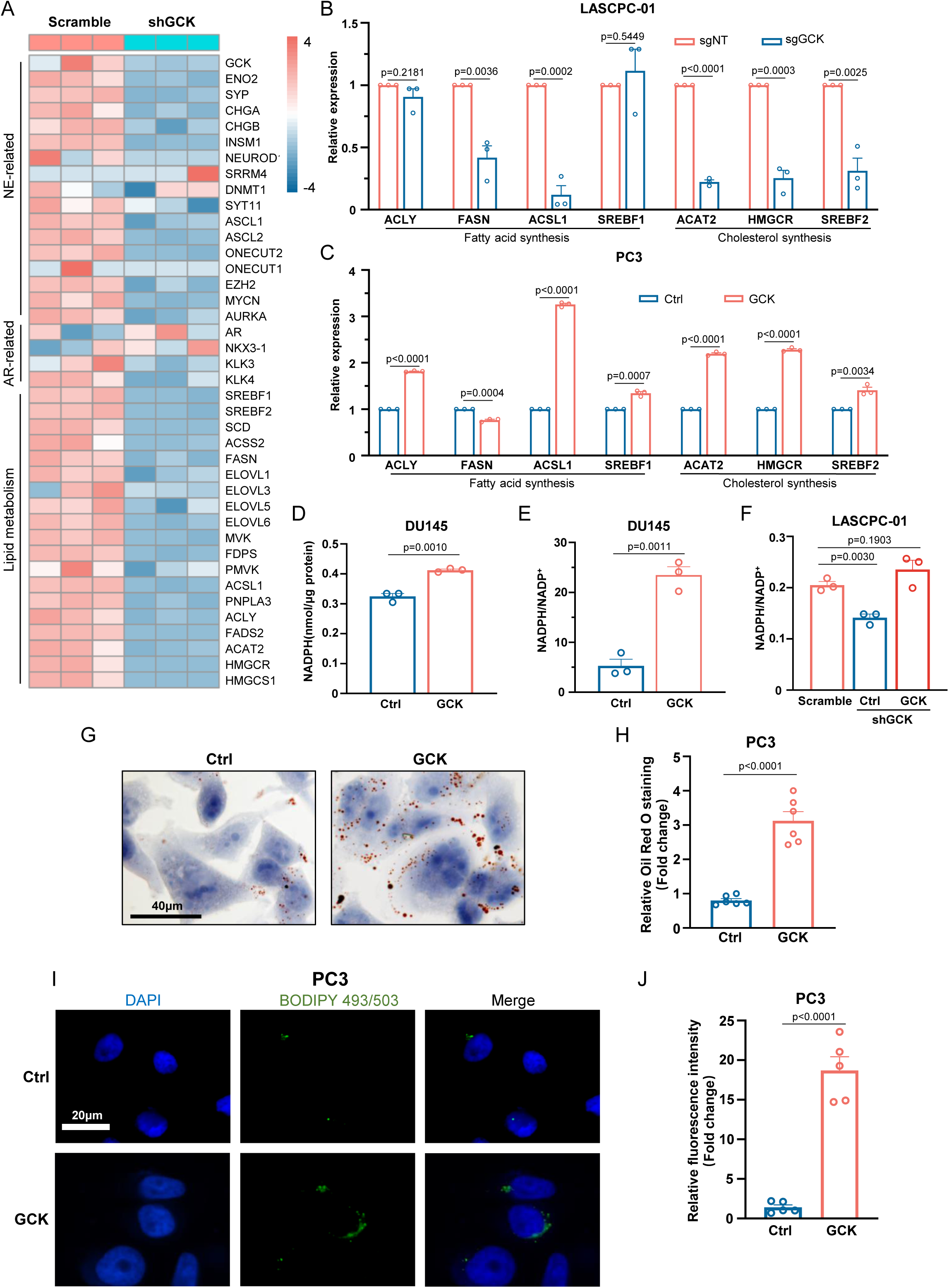
GCK promotes lipid metabolism in NEPC. (A) Heatmap showing the expression of neuroendocrine-related and lipid metabolism-related genes in shGCK and scramble of LASCPC-01 cells. (B) QPCR showing relative mRNA expression of indicated genes in LASCPC-01/sgNT and LASCPC-01/sgGCK cells (n = 3 biologically independent experiments). (C) QPCR showing relative mRNA expression of indicated genes in PC3/Ctrl and PC3/GCK cells (n = 3 biologically independent experiments). (D) Total NADPH levels in DU145/Ctrl and DU145/GCK cells (n = 3 biologically independent experiments). (E) NADPH/NADP^+^ ratio levels in DU145/Ctrl and DU145/GCK cells. (F) NADPH/NADP^+^ ratio levels in indicated LASCPC-01 cells. (G) Representative images of Oil Red O staining in PC3/Ctrl and PC3/GCK. Scale bar, 40 μm. (H) Quantification of Oil Red O staining intensity in PC3/Ctrl and PC3/GCK (n = 6 biologically independent experiments). (I) Representative immunofluorescence staining of BODIPY 493/503 and DAPI in PC3/Ctrl and PC3/GCK. Scale bar, 20 μm. (J) Quantification of relative fluorescence intensity of BODIPY 493/503 in PC3/Ctrl and PC3/GCK (n = 6 biologically independent experiments). Data presented as mean ± s.e.m (B, C, D, E, F, H, J). Statistical significance was determined by two-tailed unpaired Student’s test (B, C, D, E, F, H, J).

**Figure S5.**
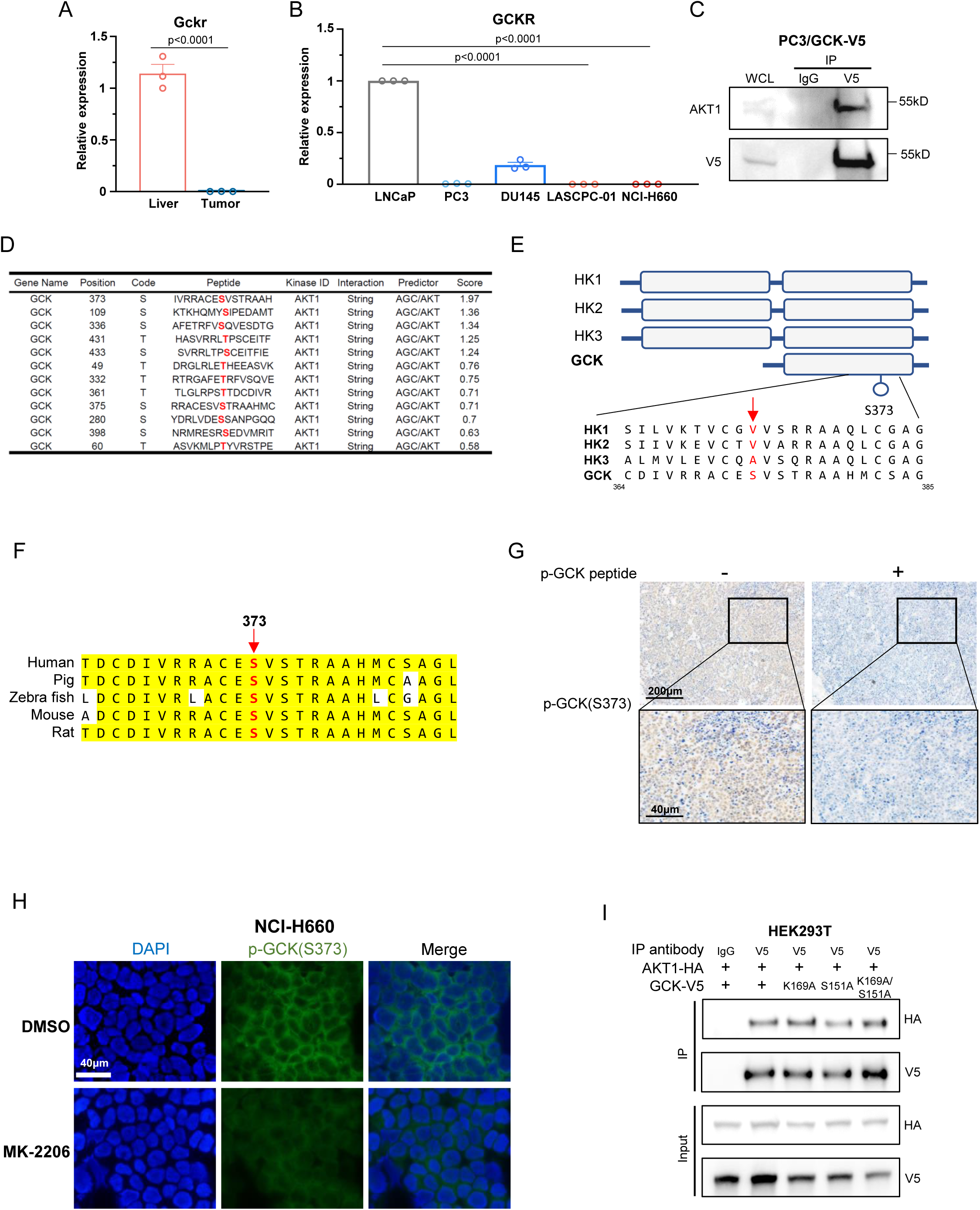
AKT1 directly binds to and phosphorylates GCK at S373. (A) QPCR showing relative mRNA expression of *Gckr* in tumor or liver tissues from Pb-Cre4: *Pten*^fl/fl^; *Trp53*^fl/fl^; *Rb1*^fl/fl^ mice (n = 3 for each group). (B) QPCR showing relative mRNA expression of *GCKR* in indicated prostate cancer cells (n = 3 biologically independent experiments). (C) Immunoprecipitation and immunoblotting assays in PC3/GCK cells (n = 3 biologically independent experiments). (D) The predicted phosphorylation sites of AKT1 on GCK using iGPS software. (E) Schematic diagram of different isoforms of hexokinase family members. (F) Alignment of protein sequences spanning GCK S373 from different species. (G) Representative immunohistochemistry staining of p-GCK(S373) in LASCPC-01 xenografts with/without recombinant p-GCK(S373) peptide treatment. Top: overview (scale bar, 200 μm); bottom: magnified view (scale bar, 40 μm). (H) Immunofluorescence images in NCI-H660 cells treated with MK-2206 (1 μM) or DMSO for 12 hours. Scale bar, 40 μm. (I) Immunoprecipitation and immunoblotting assays in HEK293T cells with indicated transfection for 48 hours (n = 3 biologically independent experiments). Data presented as mean ± s.e.m (A, B). Statistical significance was determined by two-tailed unpaired Student’s test (A, B).

**Figure S6.**
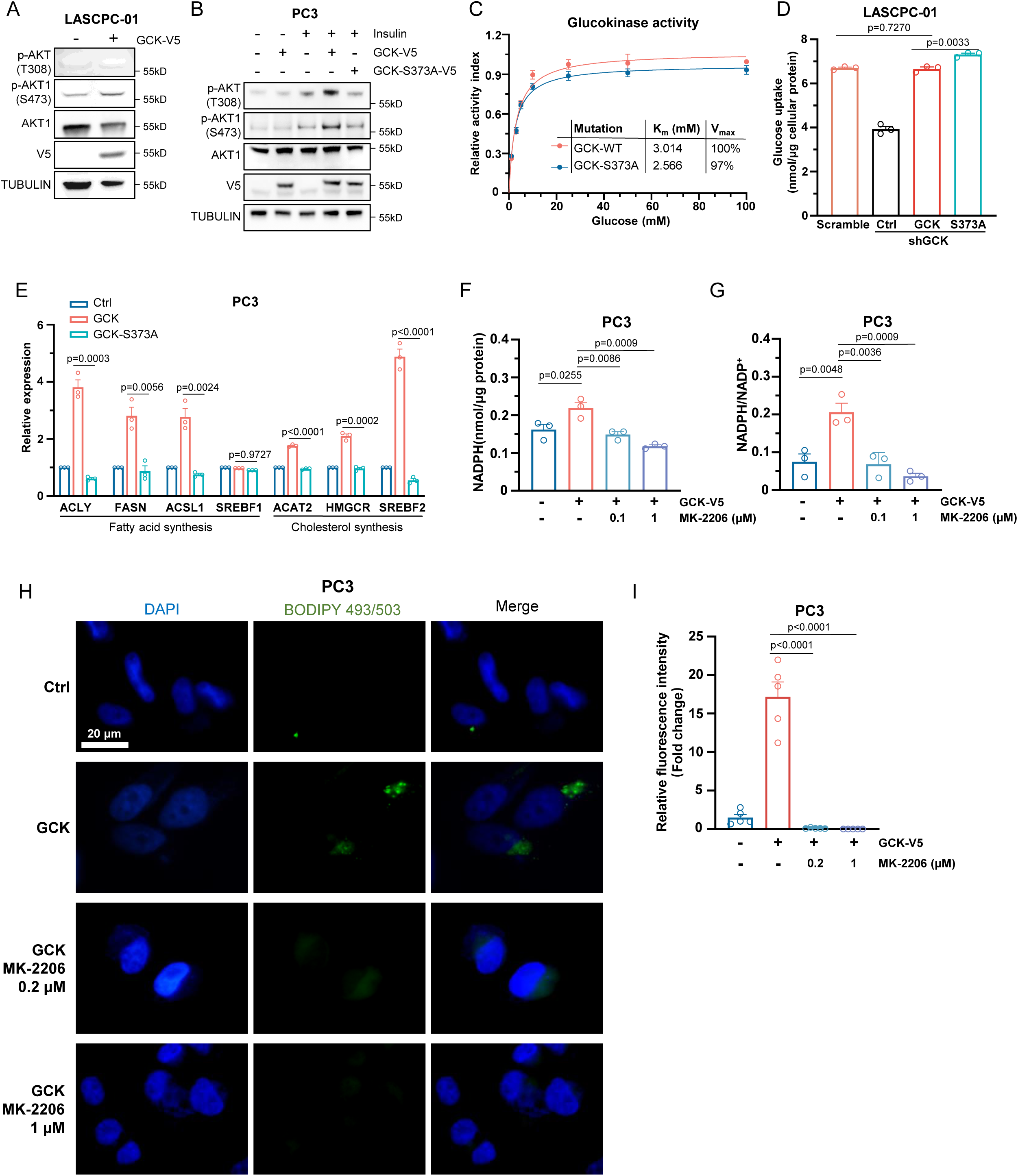
GCK functions as a protein kinase and phosphorylates AKT1. (A) Western blot showing indicated protein expression in LASCPC-01/Ctrl and LASCPC-01/GCK (n = 3 biologically independent experiments). (B) Western blot showing indicated protein expression in PC3/Ctrl, PC3/GCK, and PC3/GCK-S373A cells treated with or without insulin (100 nM) for 20 minutes (n = 3 biologically independent experiments). (C) Glucokinase activity assay in GCK-WT-V5 and GCK-S373A-V5 purified from HEK293T cell lysate with V5 antibody (n = 3 biologically independent experiments). (D) Glucose uptake assay in indicated LASCPC-01 cells (n = 3 biologically independent experiments). (E) QPCR showing relative mRNA expression of indicated genes in PC3/Ctrl, PC3/GCK, and PC3/GCK-S373A (n = 3 biologically independent experiments). (F) Total NADPH levels in PC3/Ctrl and PC3/GCK treated with the indicated concentrations of MK-2206 for 12 hours (n = 3 biologically independent experiments). (G) NADPH/NADP^+^ ratio levels in PC3/Ctrl and PC3/GCK treated with the indicated concentrations of MK-2206 for 12 hours. (H) Representative immunofluorescence staining of BODIPY 493/503 and DAPI in PC3/Ctrl and PC3/GCK treated with the indicated concentrations of MK-2206 for 12h. Scale bar, 20 μm (I) Quantification of relative fluorescence intensity of BODIPY 493/503 in PC3/Ctrl and PC3/GCK with indicated treatment (n = 5 biologically independent experiments). Data presented as mean ± s.e.m (C, D, E, F, G, I). Statistical significance was determined by two-tailed unpaired Student’s test (D, E, F, G, I) or one-way analysis of variance (ANOVA) with Dunnett’s multiple comparisons (C).

**Figure S7.**
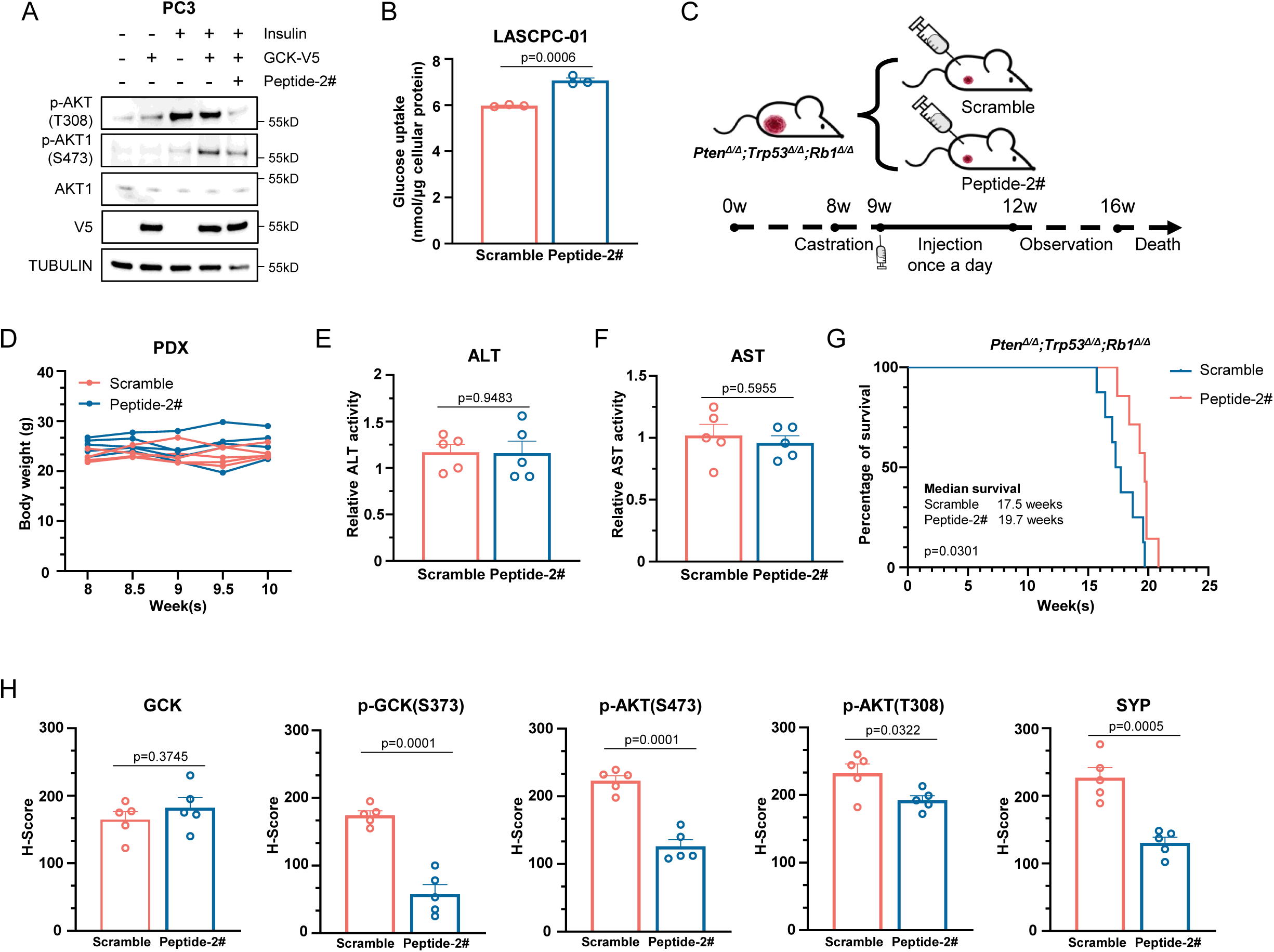
Targeting phosphorylation of GCK S373 via a peptidic inhibitor blocks lineage reprogramming. (A) Western blot showing protein expression of AKT1, p-AKT(T308), and p-AKT1(S473) in PC3/Ctrl and PC3/GCK cells with indicated treatment. (B) Glucose uptake assay in LASCPC-01 treated with scramble or peptide 2# (n = 3 biologically independent experiments). (C) Schematic illustration of Pb-Cre4: *Pten*^fl/fl^; *Trp53*^fl/fl^; *Rb1*^fl/fl^ mice treated with scramble or peptide 2# (n = 7 per group). (D) Body weight changes of NEPC PDX nude mice treated with scramble or peptide 2#. Body weight was measured twice a week at indicated time points (n = 5 per group). (E) Relative ALT activity in NEPC PDX nude mice treated with scramble or peptide 2# (n = 5 per group). (F) Relative AST activity in NEPC PDX nude mice treated with scramble or peptide 2# (n = 5 per group). (G) Kaplan-Meier survival analysis of Pb-Cre4: *Pten*^fl/fl^; *Trp53*^fl/fl^; *Rb1*^fl/fl^ mice treated with scramble or peptide 2# (n = 7 per group). (H) Quantification of GCK, p-GCK (S373), p-AKT (S473), p-AKT (T308), and SYP staining in NEPC PDX treated with scramble or peptide 2#, each dot represents sample from an individual mouse (n = 5 per group). Data presented as mean ± s.e.m (B, D, E, H). Statistical significance was determined by two-tailed unpaired Student’s test (B, D, E, H) or Mantel–Cox test (G).

